# Mitochondrial pyruvate supports lymphoma proliferation by fueling a non-canonical glutamine metabolism pathway

**DOI:** 10.1101/2021.08.26.457847

**Authors:** Peng Wei, Alex J. Bott, Ahmad A. Cluntun, Jeffrey T. Morgan, Corey N. Cunningham, John C. Schell, Yeyun Ouyang, Scott B. Ficarro, Jarrod A. Marto, Nika N. Danial, Ralph J. DeBerardinis, Jared Rutter

**Author notes:** Correspondence (J. Rutter).

## Abstract

The fate of pyruvate, which is modulated by the activity of the mitochondrial pyruvate carrier (MPC), is a defining metabolic feature in many cancers. Diffuse large B-cell lymphomas (DLBCLs) are a genetically and metabolically heterogeneous cancer. Although MPC expression and activity differed between DLBCL subgroups, mitochondrial pyruvate was uniformly consumed by glutamate pyruvate transaminase 2 (GPT2) to support α-ketoglutarate production as part of glutaminolysis. This led us to discover that glutamine exceeds pyruvate as a carbon source for the tricarboxylic acid (TCA) cycle in DLBCLs. Furthermore, we found that MPC inhibition unexpectedly leads to decreased glutaminolysis, which is contrary to previous observations in other cell types. We also discovered that MPC inhibition and depletion only decreased DLBCL proliferation in an extracellular matrix (ECM) environment and *in vivo* xenografts, but not in the typical DLBCL suspension environment. We also have found that the metabolic profile of DLBCL cells in ECM is markedly different from cells in suspension environment. Thus, we report that besides the canonical glutamate dehydrogenase (GDH)-mediated glutaminolysis, the non-canonical GPT2 mediated consumption and assimilation of glutamine and pyruvate in DLBCLs enables their proliferation in an extracellular environment-dependent manner.

**HIGHLIGHTS:**

- Glutamine, but not glucose, is a major carbon source for the tricarboxylic acid cycle in diffuse large B-cell lymphomas.
- Mitochondrial pyruvate supports glutaminolysis in diffuse large B-cell lymphomas by supplying pyruvate for glutamate pyruvate transaminase 2-mediated α -ketoglutarate production.
- Mitochondrial pyruvate carrier inhibition leads to decreased glutaminolysis in diffuse large B-cell lymphomas.
- α -ketoglutarate production is important for diffuse large B-cell lymphoma proliferation in a solid extracellular matrix environment.
- Mitochondrial pyruvate carrier activity supports diffuse large B-cell lymphoma proliferation in a solid extracellular matrix environment and in mouse xenografts.

## INTRODUCTION

The central pathway of carbohydrate metabolism is the conversion of glucose to pyruvate via glycolysis in the cytosol. The subsequent fate of this pyruvate is a critical metabolic node in mammalian cells. In most differentiated cells, the mitochondrial pyruvate carrier (MPC) transports pyruvate into mitochondria where it is used to fuel oxidation and anabolic reactions (Bricker et al., 2012). In contrast, in stem cells and many cancers, pyruvate is primarily converted to lactate and excreted from the cell (Vander Heiden et al., 2009), a process known as the Warburg effect. Several groups have shown in a variety of tumor types that the Warburg effect can be caused by low activity of the MPC (Schell et al., 2014; Li et al., 2017; Tang et al., 2019; Zou et al., 2019). Moreover, re-expression of the MPC in colon cancer cells, which have very low native expression, not only increased mitochondrial pyruvate oxidation but also repressed tumor growth (Schell et al., 2014). However, repression of the MPC is not a universal feature of all cancers. Indeed, in prostate cancer, high MPC activity is required for lipogenesis and oxidative phosphorylation, and, in hepatocellular carcinoma, high MPC activity is required to supply mitochondrial pyruvate for the tricarboxylic acid (TCA) cycle (Bader et al., 2019; Tompkins et al., 2019). Therefore, we lack a unified understanding of the relationship between a given cancer type and its dependence on the MPC and mitochondrial pyruvate.

Diffuse large B-cell lymphomas (DLBCLs) are the most common type of non-Hodgkin lymphoma and are genetically and phenotypically heterogeneous (Abramson and Shipp, 2005; Lenz and Staudt, 2010). This genetic heterogeneity has been captured by independent classification schemes (Alizadeh et al., 2000; Monti, 2005). In particular, consensus cluster classification has identified three subgroups of DLBCL based on gene expression and metabolic signatures: BCR-DLBCL, which are characterized by the expression of genes encoding B-cell receptor (BCR) signaling pathways; OxPhos-DLBCL, which have high expression of genes involved in mitochondrial oxidative phosphorylation; and the HR-DLBCL, which have increased expression of genes involved in host inflammatory infiltration (Monti, 2005; Caro et al., 2012; Norberg et al., 2017).

In terms of metabolism, OxPhos-DLBCLs display greater fatty acid oxidation than BCR-DLBCLs, whereas BCR-DLBCLs have a higher rate of glycolysis (Caro et al., 2012). However, the qualitative and quantitative differences of carbohydrate metabolism and the broader spectrum of metabolic substrates feeding the TCA cycle, between Oxphos-DLBCLs and BCR-DLBCLs, including glutaminolysis and the interplay between different fuels, is not fully understood. Understanding these basic features of metabolism in these two DLBCL subgroups might inform their therapeutic vulnerabilities.

Although DLBCLs primarily form solid tumors (Bakhshi and Georgel, 2020; Chiche et al., 2019), they are routinely passaged and studied in suspension media in the laboratory. Mimicry of the native architecture of a solid extracellular matrix (ECM) tumor microenvironment could be a key experimental factor in recapitulating DLBCL biology *ex vivo*. Therefore, we set out to examine the metabolic features of OxPhos-DLBCLs and BCR-DLBCLs in both suspension and Matrigel-based ECM growth conditions.

## RESULTS

### OxPhos-DLBCLs have higher MPC expression and activity than BCR-DLBCLs

OxPhos-DLBCLs exhibit elevated expression of numerous genes encoding components of the mitochondrial electron transport chain (ETC) compared with BCR-DLBCLs (Monti, 2005). Since MPC expression typically correlates with an oxidative metabolic phenotype, we hypothesized that OxPhos-DLBCLs would have higher levels of the MPC than BCR-DLBCLs. Using patient tumor microarray data (GSE10846), we found that the mRNA levels of *MPC1* and *MPC2*, the genes that encode the MPC1 and MPC2 subunits of the MPC, were indeed higher in OxPhos-DLBCLs (**Fig. 1A**). Furthermore, across ten DLBCL cell lines, we found that OxPhos-DLBCLs generally had higher MPC1 and MPC2 protein levels than BCR-DLBCLs (**Fig. 1B**). Analysis of proteomics from isolated mitochondria (Norberg et al., 2017), showed that MPC2 was 7-fold more abundant in OxPhos-DLBCLs than BCR-DLBCLs (MPC1-derived peptides were not detected) (**Fig. S1A, S1B**). Because MPC1 and MPC2 form an obligate heterodimer and their protein abundances are typically tightly linked, these data suggest that the MPC complex as a whole is upregulated in OxPhos-DLBCLs (Schell et al., 2014).

**Figure 1.**
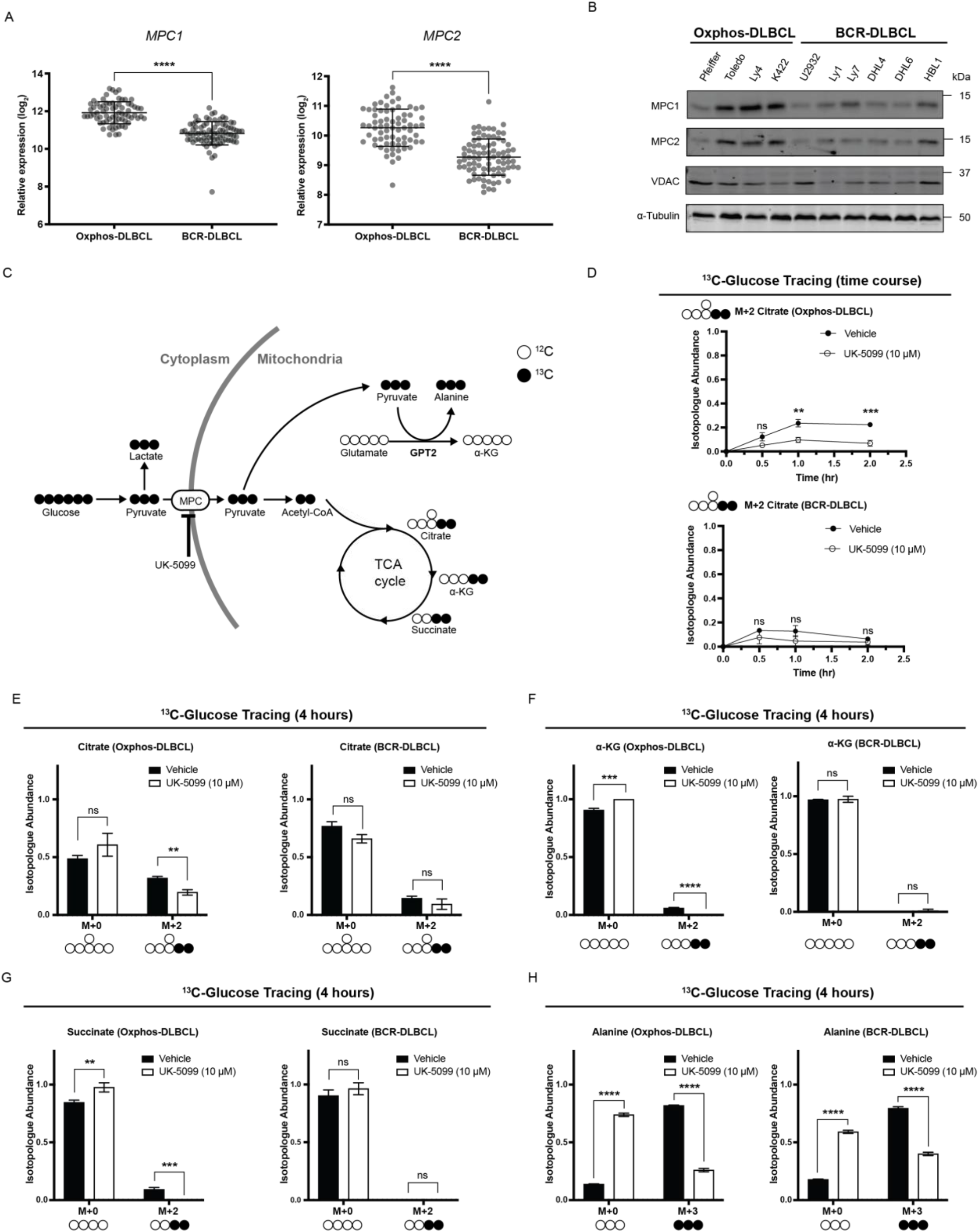
OxPhos- and BCR-DLBCLs have similar pyruvate metabolic profiles. (A) Aggregated MPC1 and MPC2 mRNA expression data from 71 OxPhos- and 83 BCR-DLBCL patient samples. Data from GSE10846. (B) Western blot analysis of MPC1, MPC2, VDAC, and α-tubulin from a panel of OxPhos- and BCR-DLBCL cell lines. (C) Schematic of D-[U-^13^C]-glucose tracing. MPC: mitochondrial pyruvate carrier. UK-5099: MPC inhibitor. α-KG: α-ketoglutarate. GPT2: glutamate pyruvate transaminase 2. (D) Quantification of the isotopologue abundance of M+2 citrate in OxPhos- and BCR-DLBCL cells cultured with D-[U-^13^C]-glucose ± the MPC inhibitor UK-5099 for 30 minutes, one hour, and two hours. Isotopologue abundance is the mean of n = 3 independent biological experiments, ± standard deviation. (E, F, G) Quantification of the isotopologue abundances of M+0 and M+2 citrate, M+0 and M+2 α-ketoglutarate (α-KG), and M+0 and M+2 succinate in OxPhos- and BCR-DLBCL cells cultured with D-[U-^13^C]-glucose ± the MPC inhibitor UK-5099 for four hours. Isotopologue abundance is the mean of n = 3 independent biological experiments, ± standard deviation. (H) Quantification of the isotopologue abundances of M+0 and M+3 alanine in OxPhos- and BCR-DLBCL cells cultured with D-[U-^13^C]-glucose ± the MPC inhibitor UK-5099 for four hours. Isotopologue abundance is the mean of n = 3 independent biological experiments, ± standard deviation. Vehicle: Dimethyl sulfoxide (DMSO) ns p >0.05; *p < 0.05; **p < 0.01; ***p < 0.001; ****p < 0.0001. Data were analyzed by one-way Anova followed by Dunnett’s multiple comparison test. See also Figure S1.

We hypothesized that the increased MPC expression in OxPhos-DLBCLs compared with BCR-DLBCLs would result in greater incorporation of carbons from glucose into the TCA cycle through increased mitochondrial transport and oxidation of pyruvate. To test this, we used D-[U-^13^C]-glucose tracing **(Fig. 1C)**. OxPhos-DLBCL (Pfeiffer) cells reached just 25% incorporation of D-[U-^13^C]-glucose into citrate by two hours, and treatment with UK-5099, a well-established MPC inhibitor (Halestrap, 1975), further decreased glucose carbon incorporation into citrate to 10% (**Fig. 1D**). BCR-DLBCL (U2932) cells reached a maximal incorporation of D-[U-^13^C]-glucose into citrate of only 15%, which was also decreased by MPC inhibition, although this change was not statistically significant (**Fig. 1D**). Thus, the difference in MPC expression levels between the two subgroups is reflected in their glucose-to-citrate labeling, with greater glucose contribution to citrate in the OxPhos subgroup–consistent with a previous study (Caro et al 2012). However, the overall contribution of glucose to citrate is minimal in both subgroups.

### DLBCLs use mitochondrial pyruvate for mitochondrial alanine synthesis

We were able to measure detectable, albeit low, levels of glucose-to-citrate labeling in DLBCL cells; however, we found even lower labeling of other TCA cycle intermediates, including α-ketoglutarate (α-KG) and succinate, from D-[U-^13^C]-glucose (**Fig. 1E, 1F, 1G; Fig. S1D**). This was especially evident in the BCR-DLBCL cell line, where very little α-KG and succinate labeling occurred even after four hours (**Fig. 1F, 1G**). Overall, this suggests that glucose does not make a substantial contribution to the TCA cycle in DLBCLs, regardless of their subtype classification. Moreover, MPC inhibition did not increase labeling of pyruvate or lactate in either OxPhos-or BCR-DLBCL cell lines (**Fig. S1C; Fig. S1D**). This contrasts with other cell types, where inhibiting mitochondrial pyruvate import leads to increased labeling of intracellular lactate, likely to compensate for the loss of ATP production from mitochondrial pyruvate oxidation (Cluntun et al., 2021). Altogether, these data suggest that pyruvate minimally contributes to mitochondrial TCA cycle metabolism and ATP production in DLBCLs.

Given the very limited degree to which carbons from glucose were incorporated into the TCA cycle, even in OxPhos-DLBCLs that exhibit higher MPC expression, we sought to uncover the destination of glucose-derived carbon following their entry into mitochondria as pyruvate. Notably, we found a striking incorporation of D-[U-^13^C]-glucose-derived carbons into alanine. This alanine labeling was dependent on MPC activity, as inhibition with UK-5099 substantially decreased labeling in both OxPhos-DLBCL and BCR-DLBCL cells (**Fig. 1H; Fig. S1C, S1D**). Thus, despite differences in MPC abundance and pyruvate oxidation in the TCA cycle, alanine is a major fate of glucose carbon in both OxPhos-DLBCLs and BCR-DLBCLs.

### Glutamine feeds the TCA cycle in an MPC-dependent manner

Alanine can be generated by the amination of pyruvate in either the mitochondria or the cytosol. Since inhibiting the MPC, and thus transport of pyruvate into the mitochondria, significantly decreased the ratio of labeled alanine in DLBCLs, our data support a model wherein alanine synthesis is predominantly mediated by the mitochondrially localized glutamate pyruvate transaminase 2 (GPT2) enzyme, which catalyzes the reversible transamination of pyruvate and glutamate to generate alanine and α-KG (**Fig. 1C**).

The GPT2 reaction and accompanying glutamate conversion to α-KG is dependent on mitochondrial pyruvate and therefore is likely dependent upon MPC activity. The robust glucose-to-alanine labeling also implies that substantial amounts of glutamine would need to be converted to mitochondrial glutamate for use by GPT2. Therefore, we tested how MPC inhibition affects glutamine consumption by DLBCLs. First, we grew DLBCLs under UK-5099 treatment for five days at a series of glutamine concentrations. The proliferation of both OxPhos- and BCR-DLBCL cells was unaffected by UK5099 at all glutamine concentrations (**Fig. 2A**), suggesting that MPC inhibition does not induce increased glutamine consumption or dependence in DLBCLs. This contrasts with the canonical metabolic responses of glioma cells, cortical neurons, and prostate cancer cells, where MPC inhibition increased glutamine consumption or dependence (Bader et al., 2019; Divakaruni et al., 2017; Yang et al., 2014).

**Figure 2.**
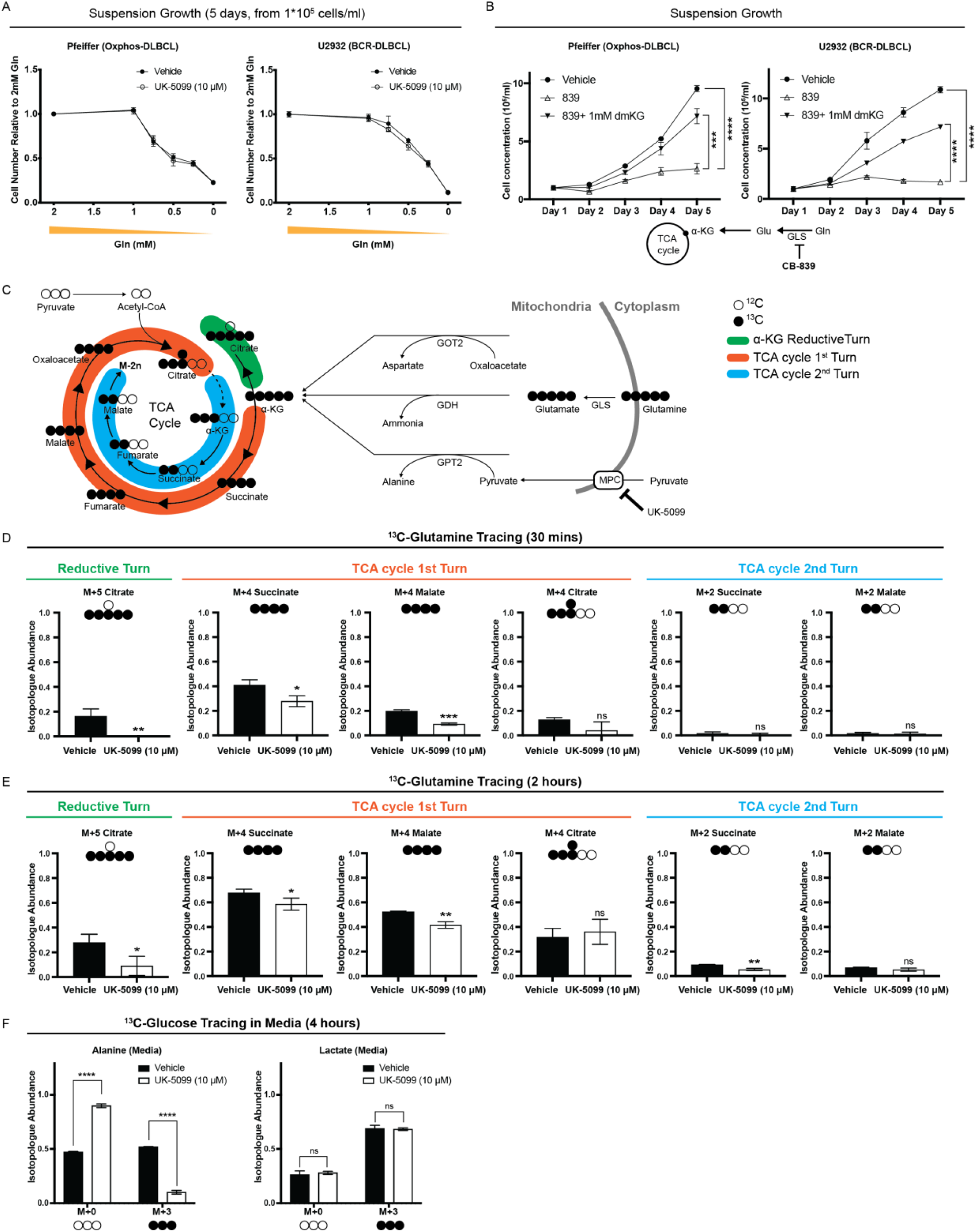
MPC inhibition affects glutamine to TCA cycle flux in DLBCL cells. (A) Growth assay of OxPhos- and BCR-DLBCL cells cultured in suspension in media supplemented with 2 mM, 1 mM, 0.75 mM, 0.5 mM, 0.25 mM, or 0 mM glutamine ± the MPC inhibitor UK-5099. Cell concentration is the mean of n = 3 independent biological experiments, ± standard deviation. (B) Growth assay of OxPhos- and BCR-DLBCL cells cultured in suspension and treated with either vehicle, GLS inhibitor CB-839, or CB-839 with dimethyl-α-ketoglutarate (dmKG). Cell concentration is the mean of n = 3 independent biological experiments, ± standard deviation. (C) Schematic of L-[U-^13^C]-glutamine tracing. α-KG: α-ketoglutarate. GOT2: mitochondrial aspartate aminotransferase. GDH: glutamate dehydrogenase. GPT2: glutamate pyruvate transaminase 2. MPC: mitochondrial pyruvate carrier. UK-5099: MPC inhibitor. (D) Quantification of the isotopologue abundances of M+5 citrate, M+4 succinate, M+4 malate, M+4 citrate, M+2 succinate, and M+2 malate in DLBCL cells cultured with L-[U-^13^C]-glutamine ± the MPC inhibitor UK-5099 for 30 minutes. Isotopologue abundance is the mean of n = 3 independent biological experiments, ± standard deviation. (E) Quantification of the isotopologue abundances of M+5 citrate, M+4 succinate, M+4 malate, M+4 citrate, M+2 succinate, and M+2 malate in DLBCL cells cultured with L-[U-^13^C]-glutamine ± the MPC inhibitor UK-5099 for two hours. Isotopologue abundance is the mean of n = 3 independent biological experiments, ± standard deviation. (F) Quantification of the isotopologue abundances of M+0 and M+3 alanine and M+0 and M+3 lactate in the medium collected from DLBCLs grown with D-[U-^13^C]-glucose ± the MPC inhibitor UK-5099 for four hours. Isotopologue abundance is the mean of n = 3 independent biological experiments, ± standard deviation). Vehicle: Dimethyl sulfoxide (DMSO) ns p >0.05; *p < 0.05; **p < 0.01; ***p < 0.001; ****p < 0.0001. Data were analyzed by one-way Anova followed by Dunnett’s multiple comparison test. See also Figure S2.

To directly test if glutaminolysis is important for DLBCL growth, we inhibited the conversion of glutamine to glutamate using the well-established glutaminase (GLS1) inhibitor CB-839 (Gross et al., 2014). Treatment with CB-839 decreased proliferation in both OxPhos- and BCR-DLBCL cells (**Fig. 2B; Fig. S2A**). Furthermore, adding a cell-permeable form of α-KG, dimethyl-α-ketoglutarate (dmKG), to DLBCLs rescued the effects of CB-839 (**Fig. 2B; S2A**), indicating that DLBCLs require α-KG generation through glutaminolysis, regardless of their subgroup classification.

To further understand how MPC inhibition affects TCA cycle metabolism, we performed a L-[U-^13^C]-glutamine isotope tracing experiment in BCR-DLBCL cells. We found that 16% of citrate exists as the M+5 isotopologue, which was completely eliminated upon MPC inhibition (**Fig. 2D**). M+5 citrate is indicative of reductive carboxylation, wherein α-KG is converted to citrate through a backwards turn of a portion of the TCA cycle (**Fig. 2C-green**) and this is thought to enable the production of citrate to fuel acetyl-CoA synthesis in the cytosol (Metallo et al., 2012; Mullen et al., 2012). Since the α-KG to citrate conversion is reversible, this result suggests the enzymes that could mediate citrate oxidation, namely isocitrate dehydrogenase 2 (IDH2) and aconitase 2 (ACO2), are active in DLBCLs.

In contrast to the minimal labeling of TCA cycle intermediates from D-[U-^13^C]-glucose, we observed substantial labeling of M+4 succinate, M+4 fumarate, M+4 malate, and M+4 citrate from L-[U-^13^C]-glutamine (**Fig. 2D; Fig. S2B**). All of these intermediates are derived from the first turn of M+5 α-KG through the TCA cycle in the oxidative direction (**Fig. 2C-orange**). Furthermore, the labeling of these TCA cycle metabolites from glutamine was significantly decreased by inhibiting the MPC (**Fig. 2D; Fig. S2B**). This contrasts with the increased glutamine anaplerosis observed in other cells upon MPC inhibition (Bader et al., 2019; Divakaruni et al., 2017; Yang et al., 2014). As expected, labeling from L-[U-^13^C]-glutamine of glutamate and various isotopomers of TCA cycle intermediates increased after two hours, but the same patterns of labeling remained evident (**Fig. 2E**). MPC inhibition again decreased this L-[U-^13^C]-glutamine labeling and the dominant isotopomers were from reductive or the first oxidative turn of the TCA cycle (**Fig. 2E; Fig. S2C**). These results suggest that DLBCLs have active glutaminolysis and α-KG oxidation and that MPC activity is required for these metabolic processes by enabling the non-canonical GPT2 mediated α-KG production.

The extensive production of alanine from glucose also suggests that alanine could play an important role in DLBCL biosynthetic processes. To address the fate of this alanine, we cultured OxPhos-DLBCL (Pfeiffer) cells with D-[U-^13^C]-glucose for four hours and collected the media for isotope tracing analysis. We found that M+3 alanine is robustly excreted from the cell and that this is dependent on MPC activity (**Fig. 2F**; **Fig. S2D**). This result supports the idea that α-KG is likely the important product of GPT2 and alanine is primarily a byproduct. We also observed M+3 lactate in the media, but, unlike M+3 alanine, MPC inhibition did not affect medium M+3 lactate abundance (**Fig. 2F**; **Fig. S2D**), similar to our previous findings for intracellular lactate labeling (**Fig. S1D**).

To summarize the above findings using L-[U-^13^C]-glutamine isotope tracing: DLBCLs have an intact and active TCA cycle, but it is primarily fed by glutamine rather than glucose. Although glucose-to-citrate labeling occurred, the glucose-derived carbon in citrate mostly did not progress through the remainder of the TCA cycle. This is likely because of citrate export to the cytosol to support biosynthesis of fatty acid and cholesterol, as well as acetylation events via acetyl-CoA synthesis (Carrer et al., 2019; Sivanand et al., 2018).

### MPC depletion reduces DLBCL proliferation in extracellular matrix and in vivo, but not in suspension environment

Given that glutamine is required as a TCA cycle fuel and for DLBCL proliferation, and mitochondrial pyruvate is required to sustain said glutamine oxidation, we hypothesized that loss of MPC function should impair proliferation of DLBCLs. However, inhibiting the MPC in cells grown in suspension culture had no effect on their proliferation (**Fig. 3A; Fig. S3A**). Because DLBCLs form solid tumors (Chiche et al., 2019), and we have previously observed that MPC-dependent effects on proliferation were particularly evident in a 3D environment (Schell et al., 2014), we investigated if MPC inhibition impairs DLBCL proliferation in Matrigel, an extracellular matrix (ECM) used to mimic the in vivo 3D environment (Benton et al., 2014). Indeed, every DLBCL cell line we tested exhibited 30–70% fewer cells when cultured with UK-5099 in growth factor reduced Matrigel for ten days (**Fig. 3B**) without any change in viability (**Fig. S3B**). These results indicate that MPC inhibition decreases the proliferation rate of DLBCLs grown in an ECM 3D environment.

**Figure 3.**
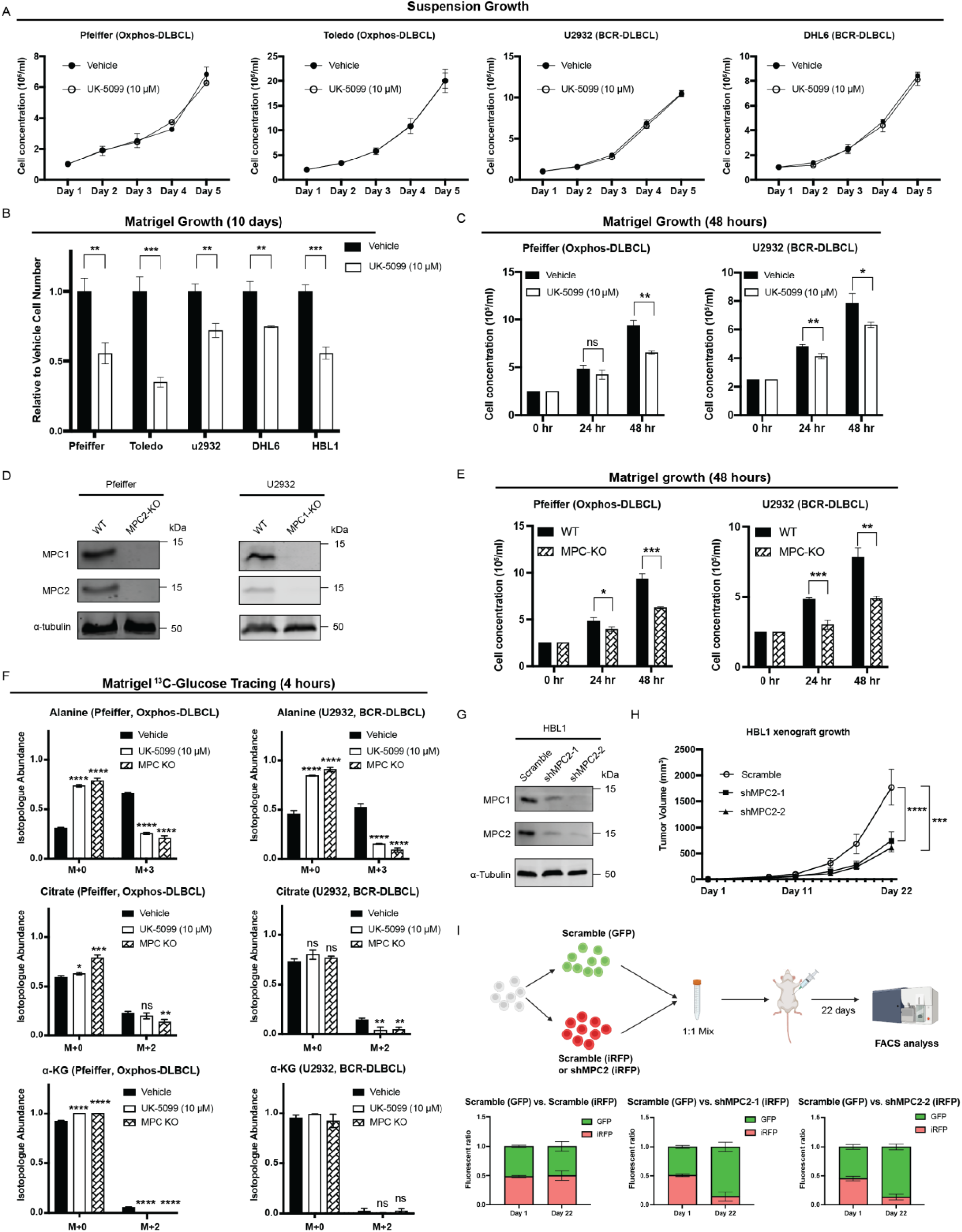
MPC inhibition reduces DLBCL proliferation in Matrigel. (A) Growth assay of DLBCL cell lines cultured in suspension ± the MPC inhibitor UK-5099 for five days. Cell concentration is the mean of n = 3 independent biological experiments, ± standard deviation. (B) Growth assay of DLBCL cell lines cultured in Matrigel ± the MPC inhibitor UK-5099 for ten days. Cell number (relative to vehicle treatment) for each cell line is the mean of n = 3 independent biological experiments, ± standard deviation. (C) Growth assay of DLBCL cell lines cultured in Matrigel ± the MPC inhibitor UK-5099 for 24 and 48 hours. Cell concentration is the mean of n = 3 independent biological experiments, ± standard deviation. (D) Western blot analysis of MPC1, MPC2, and α-tubulin in wild type (WT) and MPC knock-out (MPC KO) DLBCL cell lines. (E) Growth assay of MPC knock-out (MPC KO) cell lines and their wild type (WT) controls in Matrigel for 24 and 48 hours. Cell concentration is the mean of n = 3 independent biological experiments, ± standard deviation). (F) Quantification of the isotopologue abundances of M+0 and M+3 alanine, M+0 and M+2 citrate, and M+0 and M+2 α-ketogluturate (α-KG) from DLBCL wild type (WT) cells or MPC knock-out (MPC KO) cells cultured in Matrigel with D-[U-^13^C]-glucose ± the MPC inhibitor UK-5099 for four hours. Isotopologue abundance is the mean of n = 3 independent biological experiments, ± standard deviation). (G) Western blot analysis of MPC1, MPC2, and α-tubulin in control (Scramble) and MPC2 knock-down (shMPC2-1 and shMPC2-2) HBL1 cell lines. (H) Xenograft tumor volume of control (Scramble) and *MPC2* knock-down (shMPC2-1 and shMPC2-2) HBL1 cell lines. Tumor volumes were determined by caliper measurement and shown as mean of n = 10 ± standard error of the mean. (I) Top, schematic of the experiment. Control (Scramble) and *MPC2* knock-down (shMPC2-1 and shMPC2-2) HBL1 cell lines were labeled with GFP (Scramble) or iRFP (Scramble or *MPC2* knock-down). GFP and iRFP labeled cells were mixed in 1:1 ratio, and subcutaneously injected into mice. The GFP:iRFP ratio of each tumor was measured 21 days after injection by flow cytometry. Bottom, the GFP:iRFP ratios of tumors from Scramble (GFP) + Scramble (iRFP), Scramble (GFP) + shMPC2-1 (iRFP), and Scramble (GFP) + shMPC2-2 (iRFP) in mice at day 1 and day 22, the ratio was shown as mean of n=3 (day 1) and n=10 (day22) ± standard deviation. Vehicle: Dimethyl sulfoxide (DMSO) ns p >0.05; *p < 0.05; **p < 0.01; ***p < 0.001; ****p < 0.0001. Cell proliferation, viability, and tracing data were analyzed by one-way Anova. Mouse Xenograft data were analyzed by two-way Anova. See also Figure S3.

When grown in the ECM, DLBCLs form compact colonies within 4–5 days of seeding. To address whether this colony formation was necessary for the MPC-dependent decrease in proliferation, we assayed DLBCL cell concentration 24 and 48 hours after plating in ECM. We found that inhibition of the MPC significantly decreased the cell concentration of all four DLBCL cell lines we tested within this timeframe (**Fig. 3C; Fig. S3C**). As before, MPC inhibition did not decrease the viability of DLBCLs in ECM at these time points (**Fig. S3D**). We next generated MPC knockout (KO) cell lines from Oxphos-DLBCL (Pfeiffer) and BCR-DLBCL (U2932) cells using CRISPR-based gene disruption (**Fig. 3D**). As before, MPC depletion had no impact on cell proliferation in the suspension environment (**Fig. S3E**), but significantly decreased their proliferation in ECM by 24 hours (**Fig. 3E**). These results show that growth in an ECM environment is sufficient to reveal an MPC-dependent growth phenotype in DLBCLs, and this phenotype is not DLBCL subtype-specific.

Given the environment-dependent effects on proliferation of MPC inhibition, we asked whether the metabolic effects that we had previously observed in UK-5099-treated suspension cells were also evident in ECM-grown cells. We performed D-[U-^13^C]-glucose tracing experiments with two DBLCL cell lines in ECM. As in suspension culture, alanine labeling was robust and largely MPC-dependent—as chemical or genetic ablation of MPC activity significantly decreased the abundance of M+3 alanine (**Fig. 3F**). We also observed minimal ^13^C-glucose labeling of TCA cycle intermediates, such as citrate and α-KG, which again was MPC-dependent (**Fig. 3F**). These results confirm that MPC inhibition and genetic ablation have similar effects on glucose metabolism in DLBCLs in both suspension and ECM environments.

Next, we tested whether the proliferation of DLBCL cells is MPC-dependent *in vivo*. Since Pfeiffer and U2932 cells did not form tumors in our xenograft assays, and we were also unable to successfully genetically eliminate *MPC1* or *MPC2* in HBL1 cells, we generated HBL1 cell lines wherein *MPC2* was knocked down by stable shRNA expression (**Fig. 3G**), as we have seen previously, depletion of either MPC subunit leads to the loss of the other (Schell et al., 2014; Bensard et al., 2020). In xenograft assays, tumors from the *MPC2*-shRNA cell lines grew much more slowly over time compared to the scrambled control (**Fig. 3H**). To more directly compare engraftment and proliferation of control and *MPC2* knockdown cells, we performed an *in vivo* competition assay with the fluorescently labelled *MPC2*-shRNA cells (iRFP) and Scrambled -shRNA control cells (GFP or iRFP). In this assay, equal numbers of GFP and iRFP labelled cells were mixed and injected into mice and, after 21 days, the tumors were collected and analyzed by flow cytometry to measure the GFP- and iRFP-positive populations. In this assay, both *MPC2*-shRNA cell lines were out-competed to by scrambled shRNA control cells (**Fig. 3I**). Taken together, these data suggest that MPC is indeed required for efficient DLBCL tumor growth *in vivo*.

### ECM environment induces DLBCL metabolic reprogramming

To understand the full metabolic impact of transitioning from suspension to a solid ECM environment, we collected BCR-DLBCL (U2932) cells for steady-state metabolomic analysis after growth in either suspension or ECM Matrigel, with or without MPC inhibition, for 4, 8, 12, or 24 hours. Through unbiased clustering of both samples and metabolites, we found that four hours was sufficient to induce robust changes in the metabolic landscape of ECM-grown cells (**Fig. 4A**). This change at four hours occured well before we observed a significant impact of MPC inhibition, which is most apparent at the 24-hour time point (**Fig. 4A**). These results indicate that the growth environment has a broad and relatively rapid impact on DLBCL metabolism. Next, we focused on how this environmental shift affects the glutamine and TCA cycle metabolic phenotypes that we previously observed using isotope tracing experiments. We detected more glutamine and less glutamate in ECM-grown cells than in suspension-grown cells (**Fig. 4B**). As for TCA-cycle metabolites, α-KG was higher in ECM relative to suspension (**Fig. 4C**), but MPC inhibition did not affect α-KG abundance in either growth environment (**Fig. 4C**). We observed decreased abundances of the remaining TCA cycle intermediates in ECM, especially fumarate and malate (**Fig. 4C**). Together, these changes result in an increased α-KG/citrate ratio in the ECM environment, which could further increase the reductive conversion of α-KG to citrate (Fendt et al., 2013). It also has been previously reported that changing from monolayer culture to spheroid growth enhanced the reductive α-KG to citrate reaction (Jiang et al., 2016). These results together demonstrate that the ECM environment significantly impacts DLBCL TCA cycle metabolism. Furthermore, MPC inhibition consistently decreased citrate and isocitrate abundance in ECM and suspension environments (**Fig. 4C**). This is likely due to a combinatorial effect of decreased mitochondrial pyruvate to both limit the minimal pyruvate oxidation and to decrease α-KG generation via GPT2. We also performed a L-[U-^13^C]-glutamine isotope tracing experiment in Pfeiffer OxPhos-DLBCL cells in both suspension and ECM environments. Similar to the previous U2932 glutamine tracing experiments performed in a suspension environment, MPC inhibition decreased the fractional labeling of the TCA cycle intermediates citrate and malate from glutamine at two hours (**Fig. 4D**). Since Oxphos-DLBCL and BCR-DLBCL subgroups uniformly depend upon MPC for ECM environment growth, we believe this metabolic reprogramming also happens in Oxphos-DLBCLs.

**Figure 4.**
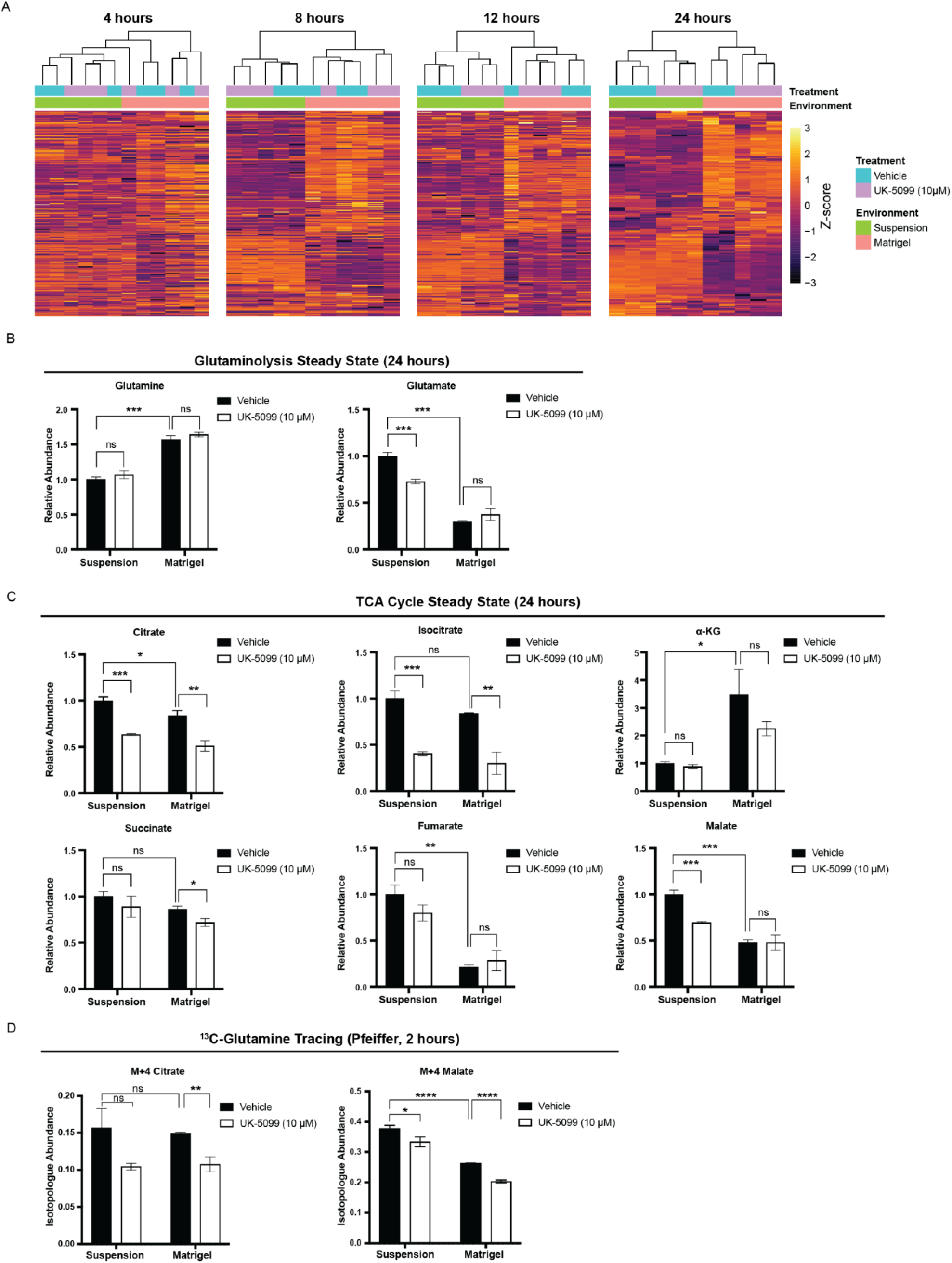
Environmental change reshapes the metabolic landscape of DLBCLs. (A) Heatmaps showing the steady-state abundances of metabolites from U2932 BCR-DLBCL cells grown either in suspension or in Matrigel ± the MPC inhibitor UK-5099 for 4, 8, 12, and 24 hours. (B) Relative steady-state abundances of glutamine and glutamate from U2932 BCR-DLBCL cells grown either in suspension or in Matrigel ± the MPC inhibitor UK-5099 for 24 hours. Metabolite abundance is the mean of n = 3 independent biological experiments, ± standard deviation. (C) Relative steady-state abundances of citrate, isocitrate, α-ketogluturate (α-KG), succinate, fumarate, and malate from U2932 BCR-DLBCL cells grown either in suspension or in Matrigel ± the MPC inhibitor UK-5099 for 24 hours. Metabolite abundance is the mean of n = 3 independent biological experiments, ± standard deviation. (D) Quantification of the isotopologue abundances of M+4 citrate and M+4 malate in Pfeiffer DLBCL cells cultured in suspension and in Matrigel with L-[U-^13^C]-glutamine ± the MPC inhibitor UK-5099 for 2 hours. Isotopologue abundance is the mean of n = 3 independent biological experiments, ± standard deviation. Vehicle: Dimethyl sulfoxide (DMSO) ns p >0.05; *p < 0.05; **p < 0.01; ***p < 0.001; ****p < 0.0001. Data were analyzed by one-way Anova followed by Dunnett’s multiple comparison test.

### DLBCLs are sensitive to ammonia in ECM

Growth in a solid ECM environment increased α-KG abundance (**Fig. 4C**), so we next asked if any of the following major α-KG-producing mitochondrial enzymes are responsible for this increase. One candidate is glutamate dehydrogenase (GDH), which converts glutamate to α-KG and produces free ammonia in the process (**Fig. 5A**). Therefore, excessive GDH activity could be toxic if free ammonia cannot be efficiently cleared (Eng et al., 2010; Kappler et al., 2017; Spanaki and Plaitakis, 2012). Furthermore, it has been reported that GDH could synthesize glutamate from α-KG and environmental ammonia, to both detoxify and recycle ammonia nitrogen for use in biosynthesis processes (Spinelli et al., 2017). A second α-KG-producing enzyme is the mitochondrial aspartate aminotransferase (GOT2), which converts glutamate to α-KG through consumption of another TCA cycle intermediate, oxaloacetate, and so does not add net carbons into the TCA cycle. (**Fig. 5A**). The third α-KG-producing enzyme, GPT2, as mentioned previously consumes glutamate and pyruvate yielding α-KG and alanine, and thus its activity is dependent upon mitochondrial pyruvate and likely MPC activity (**Fig. 5A**). Accordingly, the relative contribution of each of these enzymes—GDH, GOT2, and GPT2—to α-KG production can be differentiated based on their consumption and production of specific metabolites.

**Figure 5.**
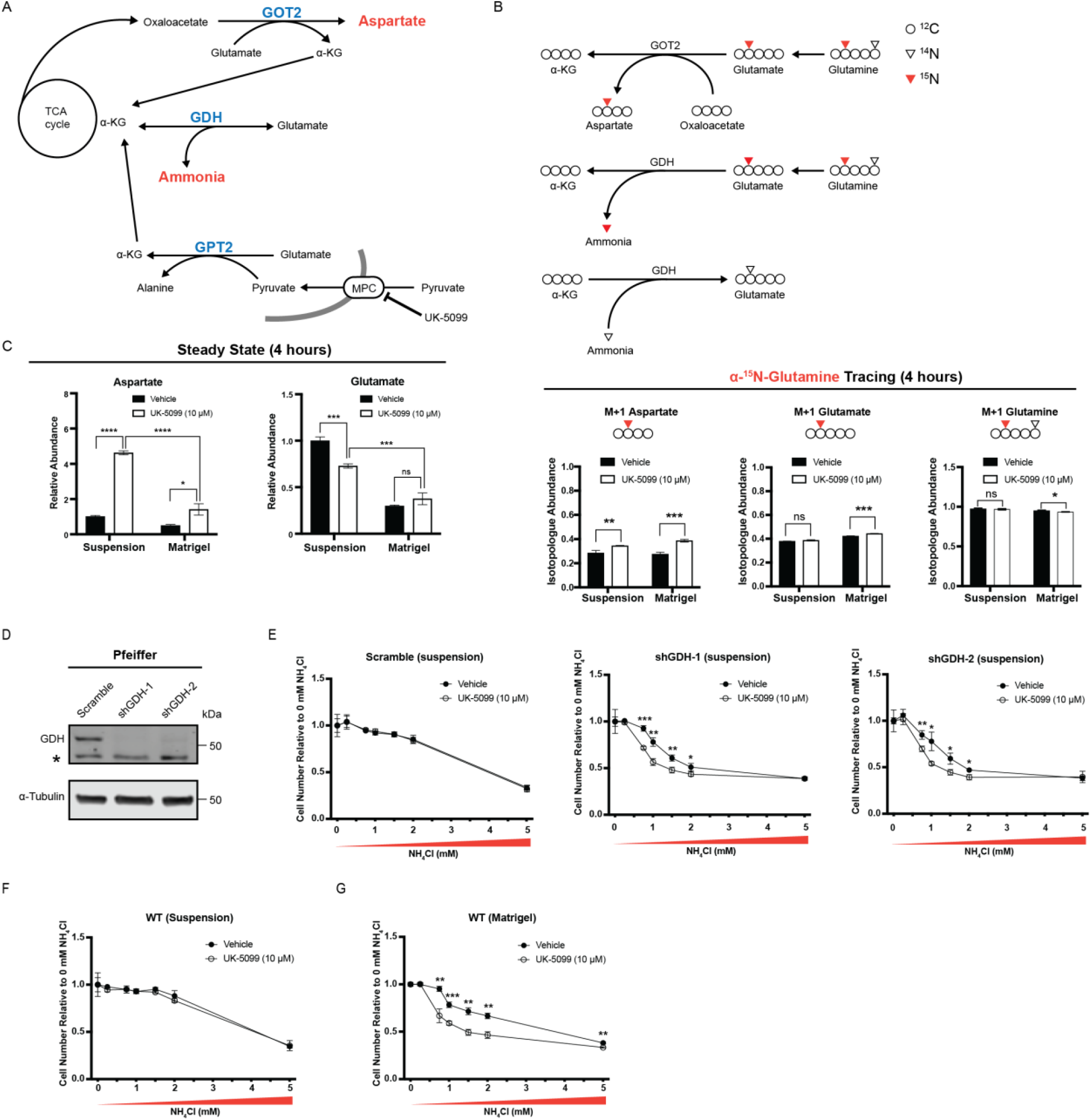
MPC inhibition enhances the sensitivity of DLBCLs to ammonia in Matrigel. (A) Schematic of the metabolic interactions of α-ketoglutarate (α-KG) production by glutamate pyruvate transaminase 2 (GPT2), mitochondrial aspartate aminotransferase (GOT2), and glutamate dehydrogenase (GDH). (B) Top, schematic of L-[alpha-^15^N]glutamine tracing. Bottom, quantification of the isotopologue abundances of M+1 aspartate, M+1 glutamate, and M+1 glutamine in DLBCL cells cultured either in suspension or in Matrigel with L-[alpha-^15^N]glutamine ± the MPC inhibitor UK-5099 for four hours. Isotopologue abundance is the mean of n = 3 independent biological experiments, ± standard deviation. (C) Relative steady-state abundance of unlabeled (^12^C) aspartate and glutamate from U2932 cells cultured either in suspension or in Matrigel ± the MPC inhibitor UK-5099 for 24 hours. Metabolite abundance is the mean of n = 3 independent biological experiments, ± standard deviation. (D) Western blot analysis of GDH and α-tubulin in control (Scramble) and *GDH* knock-down (shGDH-1 and shGDH-2) Pfeiffer DLBCL cell lines. * Indicates a non-specific band. (E) Growth assay of control (Scramble) and *GPT2* knock-down (shGDH-1 or shGDH-2) Pfeiffer DLBCL cells cultured in suspension with 0, 0.3, 0.75, 1, 1.5, 2, or 5 mM of NH_4_Cl ± the MPC inhibitor UK-5099 for 48 hours. Cell number (relative to 0 mM NH_4_Cl without UK-5099 treatment) is the mean of n = 3 independent biological experiments, ± standard deviation. (F) Growth assay of Pfeiffer cells cultured in suspension with 0, 0.3, 0.75, 1, 1.5, 2, or 5mM of NH_4_Cl ± the MPC inhibitor UK-5099 for 48 hours. Cell number (relative to 0 mM NH_4_Cl without UK-5099 treatment) is the mean of n = 3 independent biological experiments, ± standard deviation. (G) Growth assay of Pfeiffer cells cultured in Matrigel with 0, 0.3, 0.75, 1, 1.5, 2, or 5mM of NH_4_Cl ± the MPC inhibitor UK-5099 for 48 hours. Cell number (relative to 0 mM NH_4_Cl without UK-5099 treatment) is the mean of n = 3 independent biological experiments, ± standard deviation. Vehicle: Dimethyl sulfoxide (DMSO) ns p >0.05; *p < 0.05; **p < 0.01; ***p < 0.001; ****p < 0.0001. Data were analyzed by one-way Anova followed by Dunnett’s multiple comparison test.

To determine if GOT2 activity is increased in response to MPC inhibition, we cultured DLBCL cells in L-[alpha-^15^N]-glutamine-containing media for four hours and analyzed incorporation of ^15^N into aspartate (**Fig. 5B**). We found that, in both suspension and ECM environments, MPC inhibition increased labeling of M+1 aspartate (**Fig. 5C; right panel**), suggesting that MPC inhibition increases GOT2 activity in both environments. Interestingly, we also found that only 42% of glutamate was labeled from L-[alpha-^15^N]-glutamine at four hours (**Fig. 5C; right panel**), but 75% of glutamate was labeled from L-[U-^13^C]-glutamine in a similar timeframe (**Fig. S2C**). This is likely due to robust ^14^N-glutamate synthesis by GDH from α-KG and environmental ^14^N-ammonia, which has been reported to occur in human breast cancer cells (Spinelli et al., 2017). Therefore, we hypothesized that this change in GOT2 activity is due to increased cellular demand for α-KG, to compensate for impaired GPT2-mediated α-KG production. In addition, this impaired α-KG production could impair GDH-mediated incorporation of free ammonia into glutamate.

Since glutamine-to-aspartate nitrogen labeling is increased upon MPC inhibition, we next questioned if the steady-state aspartate abundance is affected by MPC inhibition. We found that in a suspension environment, MPC inhibition increased aspartate abundance by 4.5-fold, while in an ECM environment, aspartate abundance was only increased by 2-fold (**Fig. 5C; left panel**). Because aspartate synthesis is directly tied to GOT2-mediated α-KG production, this result suggests that GOT2-mediated α-KG production might be lower in ECM, perhaps due to decreased availability of oxaloacetate. Therefore, we next asked if glutamate synthesis via GDH is also affected in ECM. We found that the ECM environment caused glutamate abundance to decrease by 70% (**Fig. 5C; left panel**). MPC inhibition decreased glutamate abundance by about 30% in the suspension environment, but glutamate abundance was not affected by MPC inhibition in ECM (**Fig. 5C; left panel**). This decreased glutamate abundance from suspension to ECM raises the possibility that either the GDH-mediated α-KG and ammonia production could have increased, or GDH-mediated ammonia recycling ability could have decreased. As a consequence of this, not enough ammonia can be recycled to glutamate by GDH, resulting in a yet additional defect in the ability of the cell to detoxify excess ammonia. To address this question, we generated Pfeiffer cell lines wherein the gene encoding *GDH* was knocked down by stable shRNA expression (**Fig. 5D**). We then asked if *GDH*-shRNA cells are more sensitive to ammonia in the suspension environment. Indeed, we observed dose-dependent NH_4_Cl toxicity in scramble shRNA control cells with MPC inhibition having no additional effect (**Fig. 5E**). In contrast, MPC inhibition caused a further sensitization NH_4_Cl toxicity in two different *GDH*-shRNA cell lines (**Fig. 5E**). We next asked if wild-type (WT) DLBCLs are more sensitive to ammonia when cultured in an ECM versus in a suspension environment. In suspension conditions, we again observed dose-dependent toxicity of NH_4_Cl with MPC inhibition having no additional effect (**Fig. 5F**). However, cells cultured in ECM were hypersensitive to NH_4_Cl, which was further exacerbated by MPC inhibition (**Fig. 5G**). Therefore, DLBCLs grown in ECM are more sensitive to ammonia than cells grown in suspension; and moreover, MPC inhibition further increases ammonia sensitivity in ECM-cultured DLBCLs.

### BCAA degradation is downregulated in MPC-inhibited DLBCLs

Because MPC inhibition decreased DLBCL proliferation in ECM, we next asked which metabolic pathways are affected by MPC inhibition in ECM. Through a metabolite set enrichment analysis on our steady-state metabolomics data, we found that after 12 or 24 hours of MPC inhibition in ECM, the most affected metabolite sets were those related to branched-chain amino acid (BCAA) degradation pathways (**Fig. 6A; Fig. S4A**). BCAA degradation includes the transamination of BCAAs such as valine, leucine, and isoleucine into their branched-chain keto acids (BCKA) i.e., alpha-ketoisovalerate (KIV), ketoisocaproate (KIC), and alpha-keto-beta-methylvalerate (KMV). In these transamination reactions, α-KG is aminated to glutamate. Therefore, our results suggest that BCAA degradation to BCKAs could be regulated by the MPC through its role in α-KG production. Although the growth environment and MPC inhibition had inconsistent effects on individual BCAAs, BCKAs were all more abundant in ECM than in suspension, and decreased by MPC inhibition (**Fig. 6B**). These results suggest that MPC inhibition decreases BCKA production presumably by limiting α-KG production.

**Figure 6.**
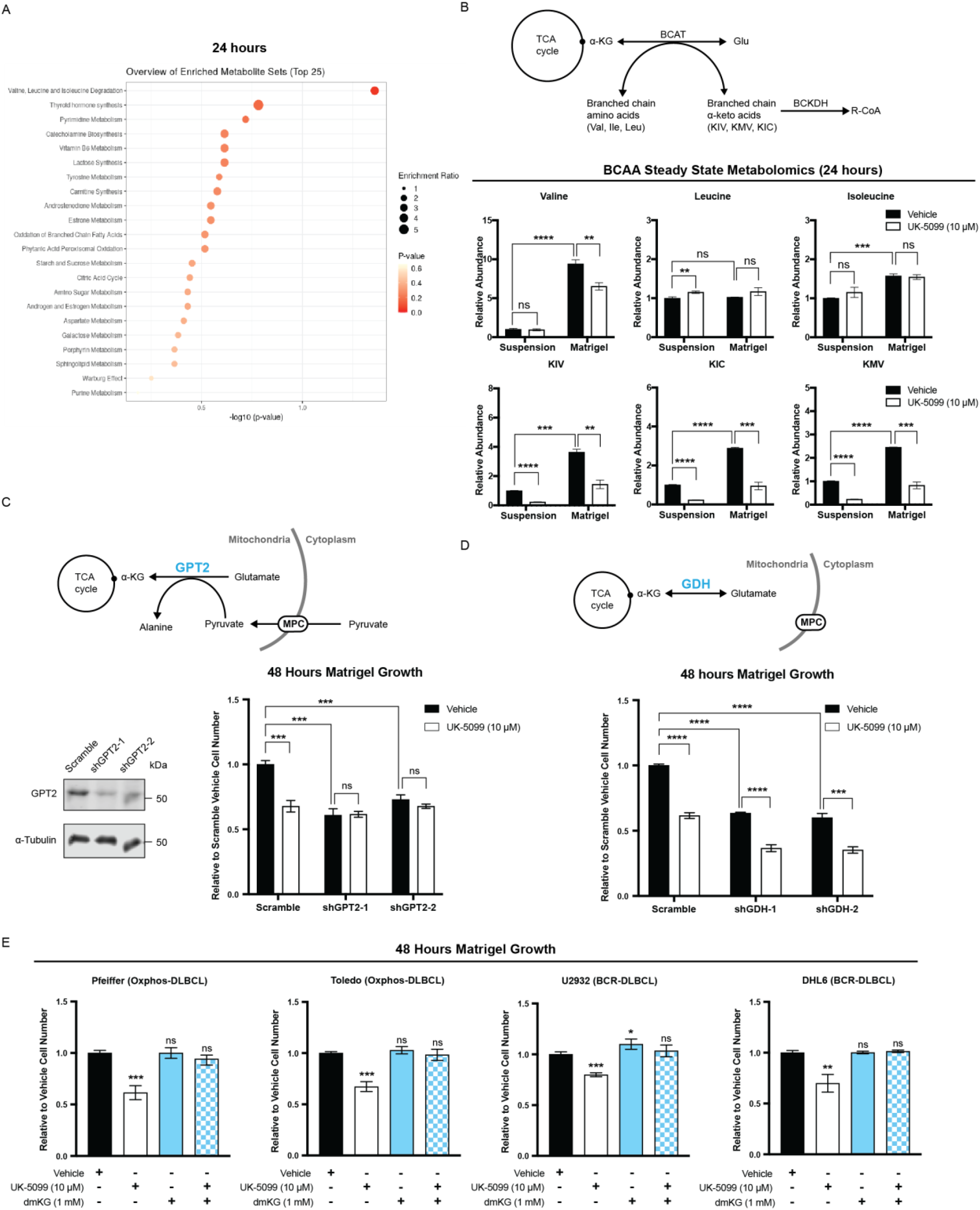
α-KG production supports DLBCL proliferation in Matrigel. (A) Metabolite set enrichment analysis based on MPC inhibition (vehicle vs. UK-5099) of DLBCL cells grown in Matrigel for 24 hours. Metabolite abundances are n = 3 independent biological experiments. (B) Top, schematic of branched-chain amino acid (BCAA) degradation pathway. Bottom, relative steady-state abundances of metabolites of the BCAA degradation pathway from DLBCL cells grown either in suspension or in Matrigel ± the MPC inhibitor UK-5099 for 24 hours. Metabolite abundance is the mean of n = 3 independent biological experiments, ± standard deviation. (C) Top, schematic of glutamate pyruvate transaminase 2 (GPT2)-mediated α-KG production pathway. Bottom left, western blot analysis of GPT2 and α-tubulin in control (Scramble) and *GPT2* knock-down (shGPT2-1 or shGPT2-2) Pfeiffer cell lines. Bottom right, growth assay of these cell lines cultured in Matrigel ± the MPC inhibitor UK-5099 for 48 hours. Cell number (relative to Scramble with vehicle treatment) is the mean of n = 3 independent biological experiments, ± standard deviation. (D) Top, schematic of GDH mediated α-KG production path. Bottom, growth assay of control (Scramble) and *GDH* knock-down (shGDH-1 or shGDH-2) Pfeiffer cell lines cultured in Matrigel ± the MPC inhibitor UK-5099 for 48 hours. Cell number (relative to Scramble with vehicle treatment) is the mean of n = 3 independent biological experiments, ± standard deviation. (E) Growth assay of DLBCL cell lines cultured in Matrigel and treated with either vehicle, UK-5099, dimethyl-α-ketoglutarate (dmKG), or UK-5099 with dmKG for 48 hours. Cell number (relative to vehicle treatment) is the mean of n = 3 independent biological experiments, ± standard deviation. Vehicle: Dimethyl sulfoxide (DMSO) ns p >0.05; *p < 0.05; **p < 0.01; ***p < 0.001; ****p < 0.0001. Data were analyzed by one-way Anova followed by Dunnett’s multiple comparison test. See also Figure S4.

### GPT2-mediated α-KG production is regulated by the MPC, and controls DLBCL proliferation

To test the hypothesis that the ECM-dependent growth defects caused by MPC inhibition were due to loss of GPT2-mediated α-KG production, we generated a *GPT2* knockdown cell line with scrambled shRNA control (**Fig. 6C**). Knockdown of *GPT2* decreased DLBCLs proliferation in ECM to a similar extent as MPC inhibition, and, importantly, we found no additive effect of MPC was inhibition (**Fig. 6C**). These results strongly support our model that MPC inhibition decreases DLBCL proliferation in ECM mainly by restricting mitochondrial pyruvate for the GPT2 reaction.

To confirm that our GPT2 findings were specific with MPC, we also tested this in *GDH* knockdown cells (**Fig. 5D**). GDH also produces α-KG, but does not require pyruvate for its reaction. Therefore, we expected *GDH* knockdown would have MPC-independent effects on DLBCL proliferation. As with *GPT2, GDH* knockdown decreased DLBCL proliferation in ECM to a similar extent as MPC inhibition (**Fig. 6D**). However, unlike *GPT2* knockdown, MPC inhibition further suppressed proliferation of *GDH* knockdown cells (**Fig. 6D**). This additive effect of MPC inhibition suggests that GDH is important for the proliferation of DLBCLs in ECM, but is likely acting through a pathway distinct from the MPC and mitochondrial pyruvate.

Finally, we added the cell-permeable form of α-KG, dmKG, to cells grown in ECM to determine if directly increasing α-KG is sufficient to rescue the effects of MPC inhibition. We found that adding dmKG alone had no effect DLBCL proliferation. However, dmKG completely rescued the MPC inhibition-dependent proliferation defect in all of the cell lines that we tested (**Fig. 6E**). This further supported the hypothesis that impaired α-KG production is the metabolic defect that underlies the MPC inhibition-induced loss of proliferation in ECM for both Oxphos- and BCR-DLBCLs.

## DISCUSSION

Based on the difference in MPC expression between OxPhos- and BCR-DLBCL subgroups, we initially set out to study potential differences in the utilization of mitochondrial pyruvate in these cell types. Indeed, OxPhos-DLBCLs display a greater incorporation of pyruvate into citrate, and are more sensitive to MPC inhibition for this incorporation metric. However, the maximum pyruvate-to-citrate labeling ratio in both OxPhos- and BCR-DLBCL subgroups is low, and labeling of TCA-cycle metabolites downstream of citrate is almost non-existent. These findings indicate that although there are differences between OxPhos- and BCR-DLBCL pyruvate metabolism, these differences are dwarfed by the effects of a common, yet unexpected, source of TCA carbon, which we discovered to be pyruvate enabled glutamine. Nevertheless, in the light of a greater understanding of the role of MPC in DLBCLs, it will be interesting to understand the differential expression of the MPC in DLBCL subtypes and whether this leads to additional phenotypes that we have yet to uncover.

Although glucose does not substantially contribute to the TCA cycle in DLBCLs, glucose-derived pyruvate facilitates the utilization of glutamine as a TCA cycle fuel by supporting GPT2-mediated α-KG production (**Fig. 7**). Glutamine tracing experiments demonstrated that glutamine-derived α-KG can be converted to citrate through either the reductive or oxidative modes of the TCA cycle (**Fig. 7A**). This suggests that DLBCLs have an intact and active TCA cycle, but we consistently observed limited oxidation of citrate to other TCA cycle intermediates. We speculate this is because citrate is being exported to the cytosol, where it can be used to produce acetyl-CoA for lipogenesis and acetylation, both of which are critical for cancer cells (**Fig. 7**) (Carrer et al., 2019; Sivanand et al., 2018). In support of this hypothesis, the mitochondrial citrate exporter SLC25A1 appears to be essential in lymphocytes and DLBCLs (**Fig. S5A, S5B**). Therefore, DLBCLs appear to have a non-canonical TCA cycle pattern that includes the export of most citrate to the cytosol. This is likely why glutamine, and not glucose, is so prominently incorporated into the TCA cycle in DLBCL cells: pyruvate-derived citrate is not used to fuel the TCA cycle, but glutamate-derived α-KG is.

**Figure 7.**
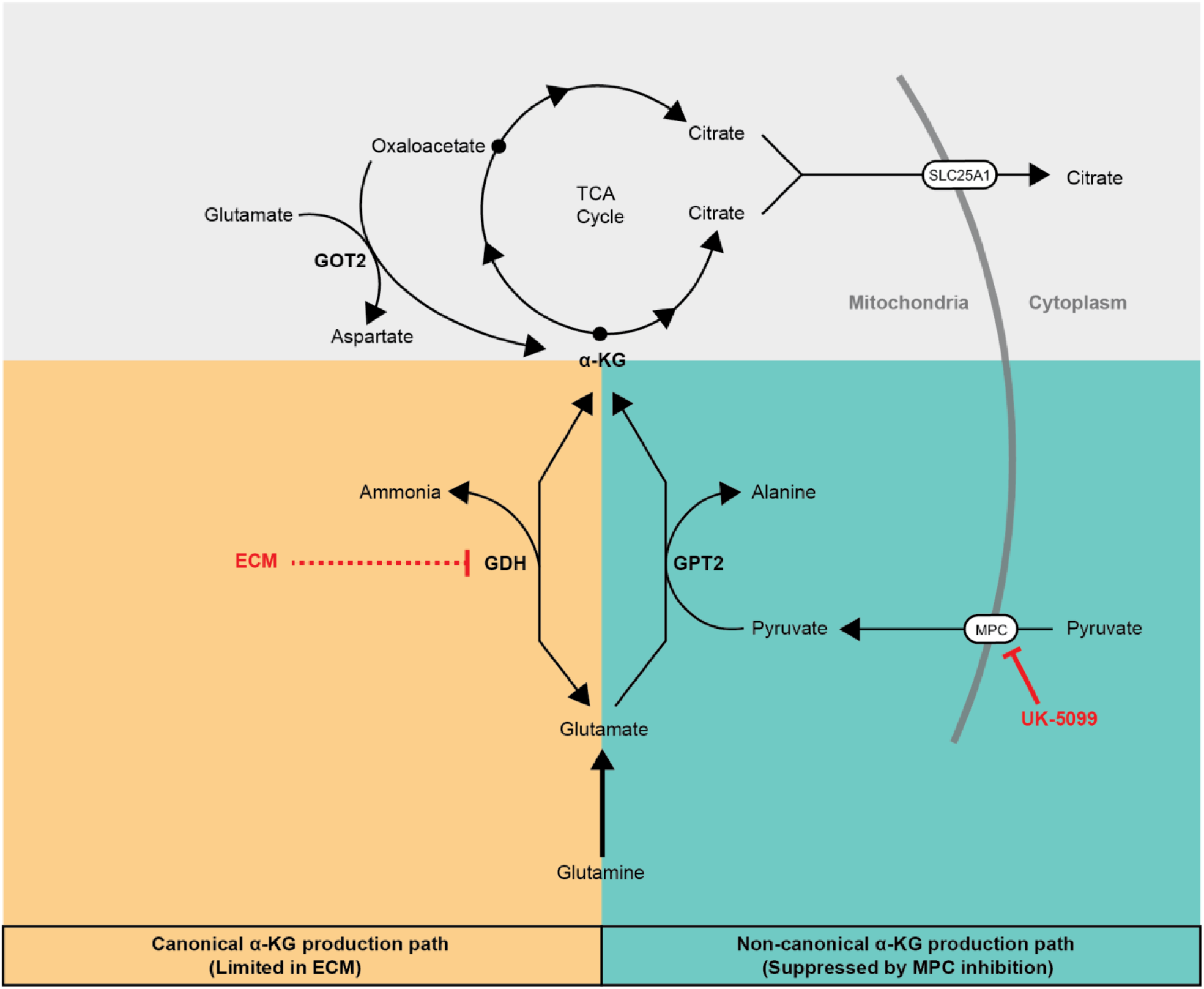
α-KG production pathways that add net carbons to TCA cycle from glutamine. α-KG: α-ketoglutarate. GOT2: mitochondrial aspartate aminotransferase. GDH: glutamate dehydrogenase. GPT2: glutamate pyruvate transaminase 2. MPC: mitochondrial pyruvate carrier. UK-5099: MPC inhibitor. ECM: extracellular matrix. See also Figure S5.

We also speculate that this previously underappreciated MPC—GPT2—α-KG axis could also play an important metabolic role outside of DLBCLs or other cancers. For example, this surprising metabolic feature of DLBCLs might be established during B-cell activation. A previous study showed that B cells increase their glucose consumption during activation, but that this increased consumption—reminiscent of our study—does not lead to labeling of TCA cycles metabolites via pyruvate (Waters et al., 2018).

Besides GPT2, GDH is another enzyme that produces α-KG from glutamate and thereby can add net carbons into the TCA cycle (**Fig. 7**). Because GDH necessarily produces free ammonia while making α-KG, high GDH activity could be toxic to the cell if free ammonia does not diffuse away and cannot be efficiently recycled. Here, we show that transitioning DLBCLs from a suspension environment to a solid Matrigel-based ECM environment makes them more sensitive to ammonia. In addition, MPC inhibition further sensitizes cells to ammonia in this solid environment. We speculate that GDH-produced ammonia does not sufficiently diffuse away from cells in a solid environment, which could feed back on the GDH reaction to prevent α-KG production. As a consequence of this decreased α-KG production, less ammonia can be recycled to glutamate by GDH, resulting in a yet additional defect in the ability of the cell to detoxify excess ammonia (**Fig. 7**).

A limitation of cell culture has always been an inability to fully recapitulate aspects of an individual cell’s organismal context. For example, the function of pyruvate metabolism for breast cancer metastasis could only be revealed in a solid growth environment (Elia et al., 2019). The importance of this limitation varies depending on cell type. It is now clear that immune cells, including B cells, function in tissues more than in the bloodstream, as previously thought (Farber, 2021). Recent studies have reported that the tissue microenvironment could influence DLBCL gene expression (Sangaletti et al., 2020), and physical properties of the extracellular matrix environment could also change cancer cells’ mitochondrial structure and function (Tharp et al., 2021). Here, we have shown that transitioning DLBCLs from a suspension environment to a solid Matrigel-based ECM rapidly reshapes their metabolome. Indeed, we found that a growth condition that better recapitulates the in vivo environment is sufficient to unveil entire facets of the DLBCL metabolic landscape not apparent in standard suspension cell culture. Indeed, we identified the environment-specific dependence on MPC activity, which we then recapitulated using in vivo tumor xenograft assays. We anticipate that other aspects of DLBCL biology are also better reflected in ECM environment, and that the metabolic requirements of many types of solid-tumor cancers may be similarly revealed by more relevant culture systems.

DLBCLs are typically more vascularized compared to follicular lymphoma (Passalidou et al., 2003; Solimando et al., 2020). Aggressive and chemotherapy-resistant DLBCLs often have high vascular endothelial growth factor expression and high microvessel density (Cardesa-Salzmann et al., 2011; Ruan and Leonard, 2009; Solimando et al., 2020). Besides importing nutrients, tumor blood vessels could also function to export metabolic byproducts, such as ammonia and lactate from the tumors. In addition, cancer cells can change their metabolism to adapt to their microenvironment, such that byproducts including ammonia and lactate become a nutrient source or anabolic substrate (Faubert et al., 2017; Spinelli et al., 2017). Since MPC inhibition affects DLBCL metabolism and induces ammonia sensitivity, it is possible that a combination therapy of an MPC inhibitor and a drug that blocks tumor blood vessel growth could sensitize cancer cells to their byproduct ammonia and also limit essential glutaminolysis.

## Acknowledgements

We thank the Preclinical Research Resource, Metabolomics Core, the Flow Cytometry Core, and the Mutation Generation & Detection Core at the University of Utah for facilitating this research, Lauren G. Zacharias and Jessica A. Sudderth in Ralph DeBerardinis’s lab for their assistance with steady-state metabolism profiling and isotope tracing experiments, and members of the Rutter lab for helpful discussion. This work was supported by 5R01CA228346 (to J.R.). A.J.B is supported by K00CA212445. J.T.M. received support as an HHMI Fellow of the Jane Coffin Childs Memorial Fund for Medical Research. C.N.C. is supported by 1F32GM140525. N.N.D. is supported by N.I.H. grant 5R01CA219850. J.R. and R.J.D. are investigators of the Howard Hughes Medical Institute. R.J.D. is supported by N.I.H. grant R35CA22044901.

## Author contributions

Conceptualization, P.W., A.J.B., and J.R.; Methodology, Formal Analysis, and Investigation, P.W., A.J.B., A.A.C., J.C.S., Y.O., S.B.F., J.A.M., N.N.D., R.J.D., and J.R.; writing – original draft, P.W.; writing – review & editing, P.W., J.T.M., C.N.C., A.J.B., J.R., A.C.C., and Y.O.; Visualization, P.W., and A.J.B.; Supervision, J.R.; Funding Acquisition, J.R.

---

**Figure S1.**
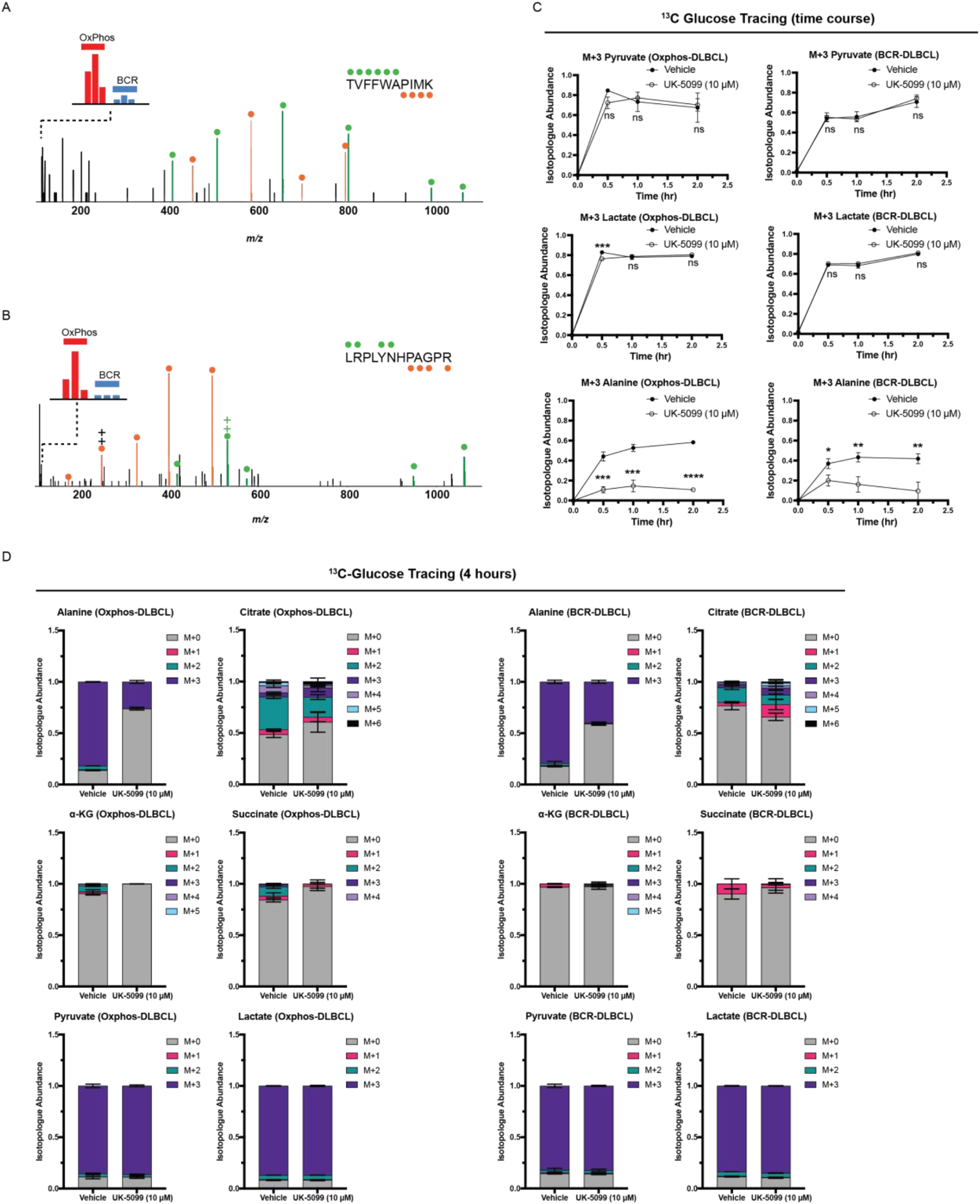
MPC expression and pyruvate metabolism in OxPhos- and BCR-DLBCLs. Related to Figure 1. (A and B) MS/MS spectra corresponding to MPC2-derived peptides 40TVFFWAPIMK49 (A) and 28LRPLYNHPAGPR39 (B) acquired during multidimensional LC-MS/MS analysis of purified mitochondria from three OxPhos- (Karpas 422, Pfeiffer, and Toledo) and three non-OxPhos/BCR- (Ly1, DHL4, and DHL6) DLBCL cell lines using DEEP SEQ mass spectrometry. Ions of b- and y-type are shown in green and orange, respectively. Relative ratios in BCR- and OxPhos-DLBCL cell lines are derived from iTRAQ reporter ion intensities shown in inset mass spectrum. (C) Quantification of the isotopologue abundance of M+2 pyruvate, M+3 lactate, and M+3 alanine in OxPhos- and BCR-DLBCL cells cultured with D-[U-^13^C]-glucose ± the MPC inhibitor UK-5099 for 30 minutes, 1 hour, and 2 hours. Isotopologue abundance is the mean of n = 3 independent biological experiments, ± standard deviation. (D) Quantification of the isotopologue abundances of alanine, citrate, α-KG, succinate, pyruvate, and lactate in OxPhos- and BCR-DLBCL cells cultured with D-[U-^13^C]-glucose ± the MPC inhibitor UK-5099 for four hours. Isotopologue abundance is the mean of n = 3 independent biological experiments, ± standard deviation. Vehicle: Dimethyl sulfoxide (DMSO) ns p >0.05; *p < 0.05; **p < 0.01; ***p < 0.001; ****p < 0.0001. Data were analyzed by one-way Anova followed by Dunnett’s multiple comparison test.

**Figure S2.**
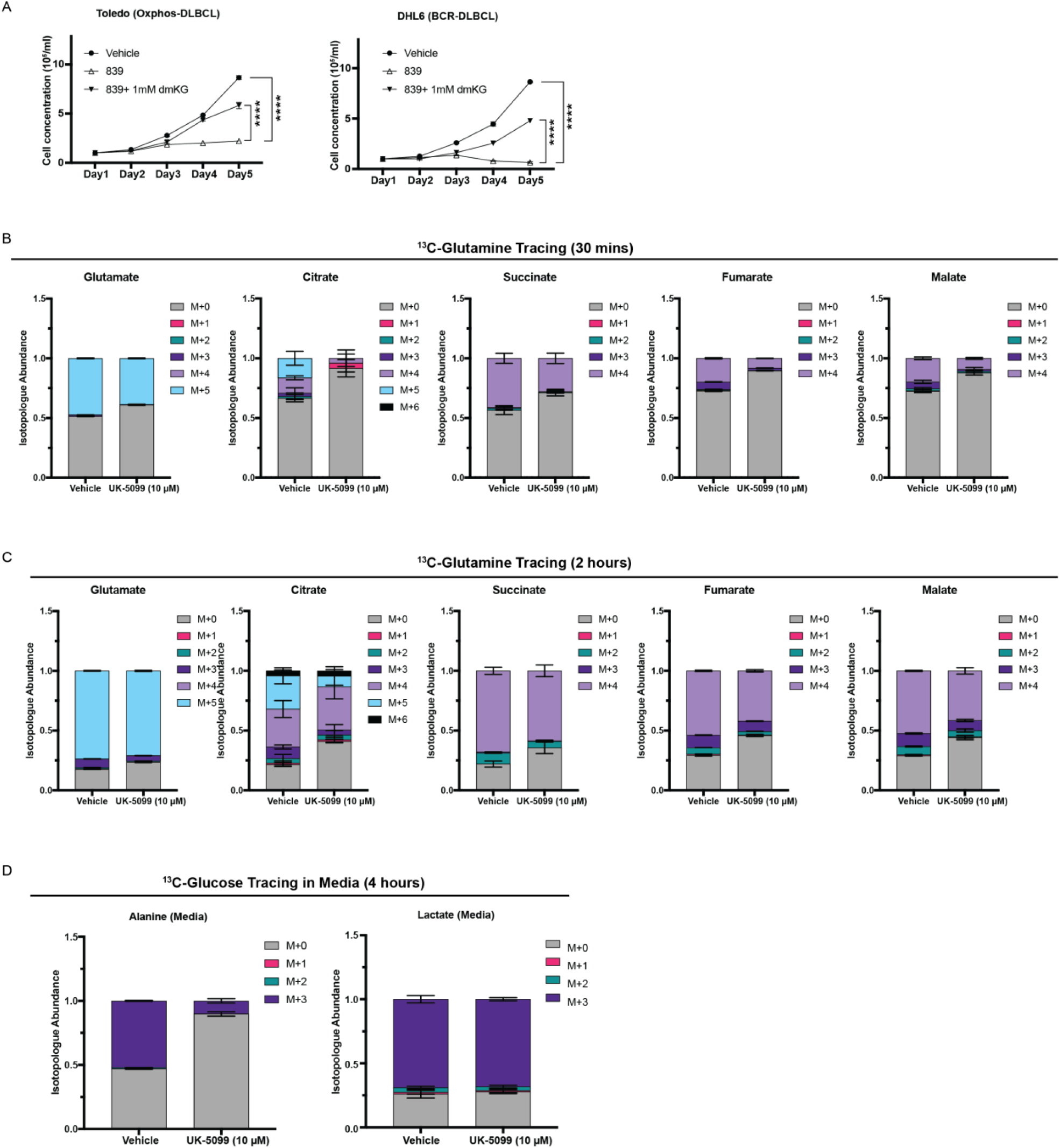
α-KG production is essential for DLBCLs proliferation, and MPC inhibition affects glutamine to TCA cycle flux in DLBCL. Related to Figure 2. (A) Growth assay of OxPhos- and BCR-DLBCL cells cultured in suspension and treated with either vehicle, GLS inhibitor CB-839, or CB-839 with dimethyl-α-ketoglutarate (dmKG). Cell concentration is the mean of n = 3 independent biological experiments, ± standard deviation. (B) Quantification of the isotopologue abundances of glutamate, citrate, succinate, fumarate, and malate in DLBCL cells cultured with L-[U-^13^C]-glutamine ± the MPC inhibitor UK-5099 for 30 minutes. Isotopologue abundance is the mean of n = 3 independent biological experiments, ± standard deviation. (C) Quantification of the isotopologue abundances of glutamate, citrate, succinate, fumarate, and malate in DLBCL cells cultured with L-[U-^13^C]-glutamine ± the MPC inhibitor UK-5099 for two hours. Isotopologue abundance is the mean of n = 3 independent biological experiments, ± standard deviation. (D) Quantification of the isotopologue abundances of alanine and lactate in the medium collected from DLBCLs grown with D-[U-^13^C]-glucose ± the MPC inhibitor UK-5099 for four hours. Isotopologue abundance is the mean of n = 3 independent biological experiments, ± standard deviation). Vehicle: Dimethyl sulfoxide (DMSO) ns p >0.05; *p < 0.05; **p < 0.01; ***p < 0.001; ****p < 0.0001. Data were analyzed by one-way Anova followed by Dunnett’s multiple comparison test.

**Figure S3.**
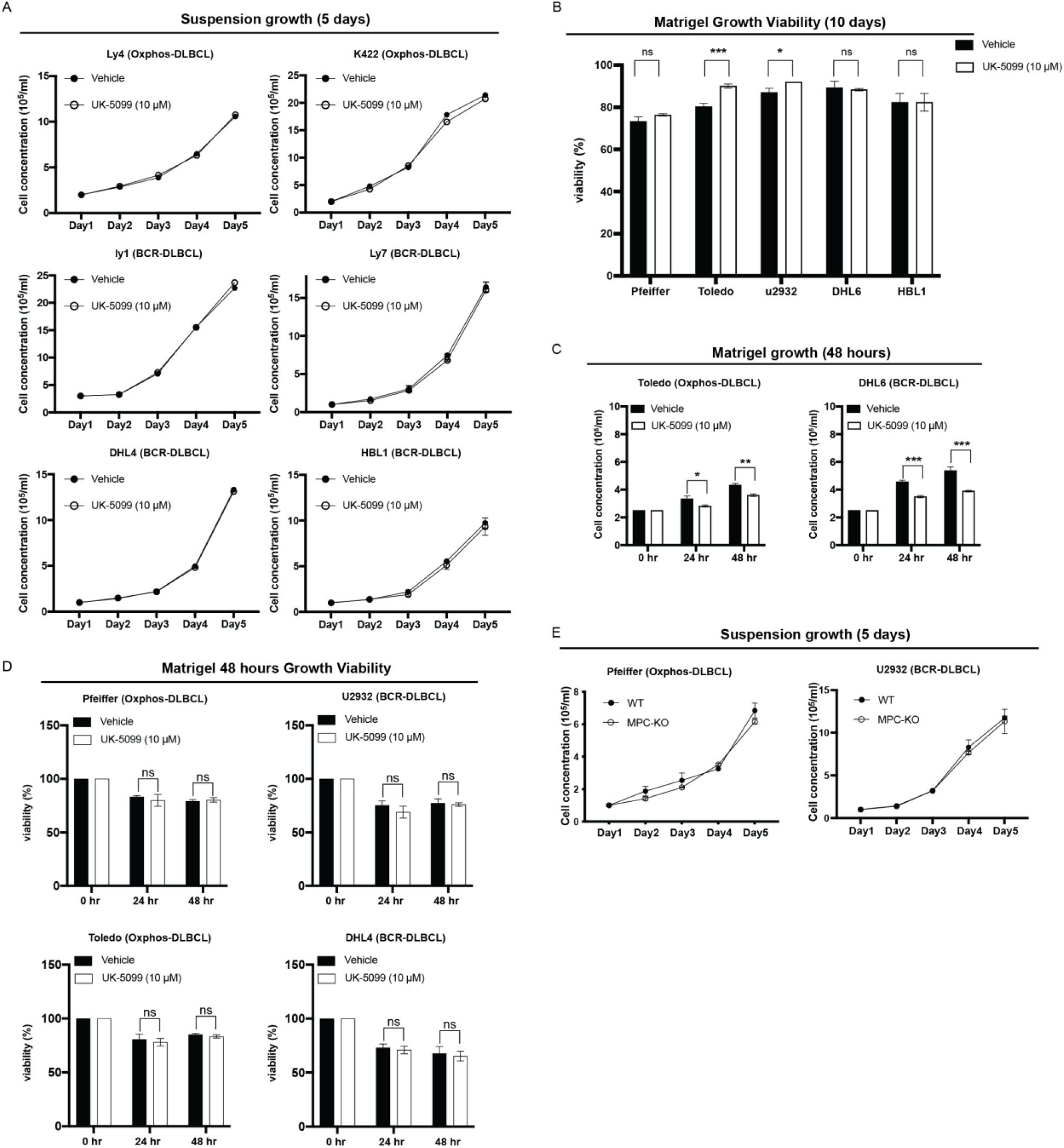
MPC inhibition reduces DLBCL proliferation in Matrigel. Related to Figure 3. (A) Growth assay of DLBCL cell lines cultured in suspension ± the MPC inhibitor UK-5099 for five days. Cell concentration is the mean of n = 3 independent biological experiments, ± standard deviation. (B) Cell viability measured by trypan blue staining of DLBCL cell lines cultured in Matrigel ± the MPC inhibitor UK-5099 for 10 days. Cell viability is the mean of n = 3 independent biological experiments, ± standard deviation. (C) Growth assay of DLBCL cell lines cultured in Matrigel ± the MPC inhibitor UK-5099 for 24 and 48 hours. Cell concentration is the mean of n = 3 independent biological experiments, ± standard deviation. (D) Cell viability measured by trypan blue staining of DLBCL cell lines cultured in Matrigel ± the MPC inhibitor UK-5099 for 48 hours. Cell viability is the mean of n = 3 independent biological experiments, ± standard deviation. (E) Growth assay of MPC knock-out (MPC-KO) cell lines cultured in suspension ± the MPC inhibitor UK-5099 for five days. Cell concentration is the mean of n = 3 independent biological experiments, ± standard deviation. Vehicle: Dimethyl sulfoxide (DMSO) ns p >0.05; *p < 0.05; **p < 0.01; ***p < 0.001; ****p < 0.0001. Data were analyzed by one-way Anova followed by Dunnett’s multiple comparison test.

**Figure S4.**
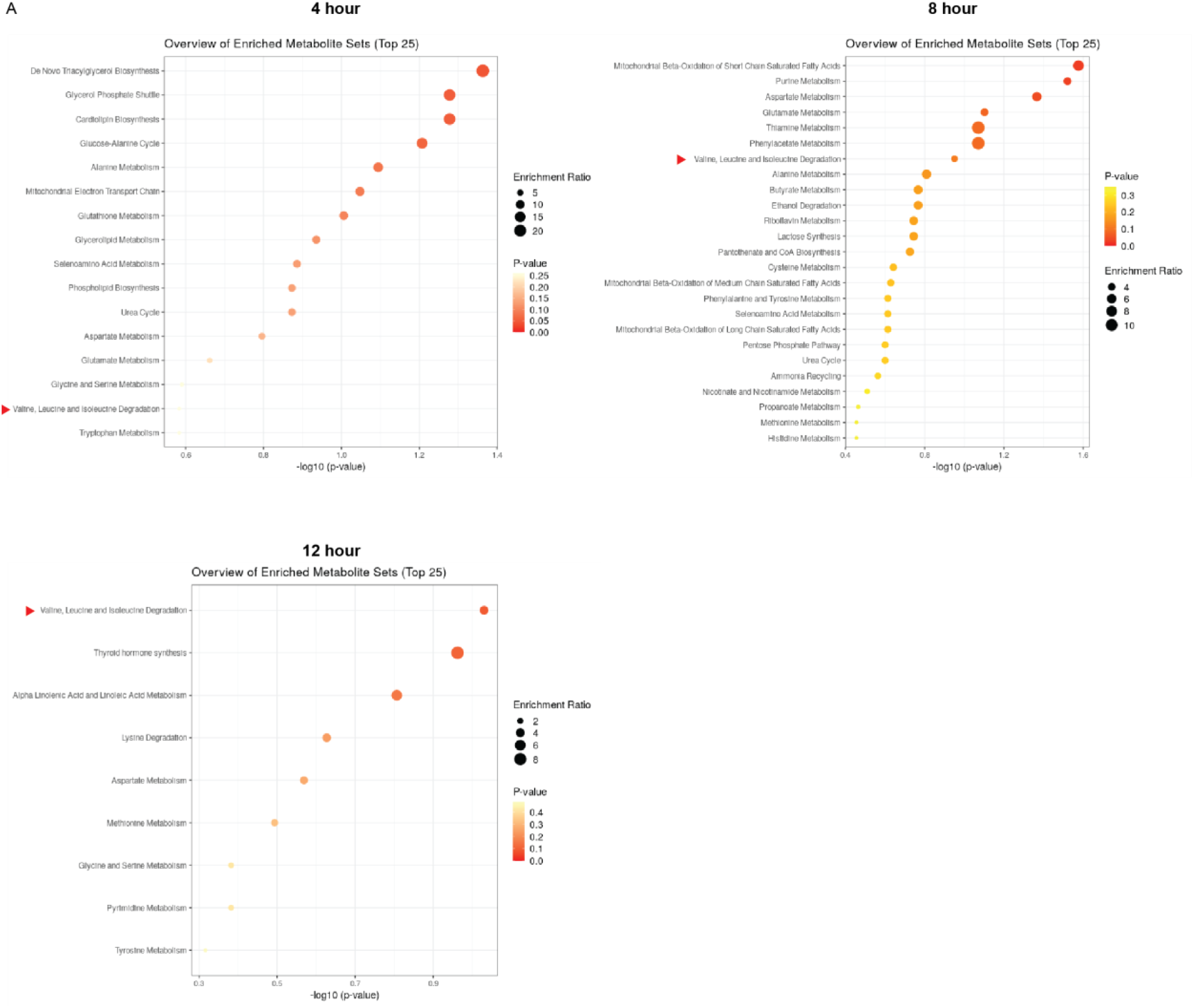
Branched-chain amino acid (BCAA) degradation pathway is affected by MPC inhibition, and α-KG is important for DLBCLs proliferation in Matrigel environment. Related to Figure 6. (A) Metabolite set enrichment analysis based on MPC inhibition (vehicle vs. UK-5099) of cells grown in Matrigel for 4, 8, and 12 hours. Metabolite abundances are n = 3 independent biological experiments. Arrowheads are pointing to BCAA degradation pathways. Vehicle: Dimethyl sulfoxide (DMSO) ns p >0.05; *p < 0.05; **p < 0.01; ***p < 0.001; ****p < 0.0001. Data were analyzed by one-way Anova followed by Dunnett’s multiple comparison test.

**Figure S5.**
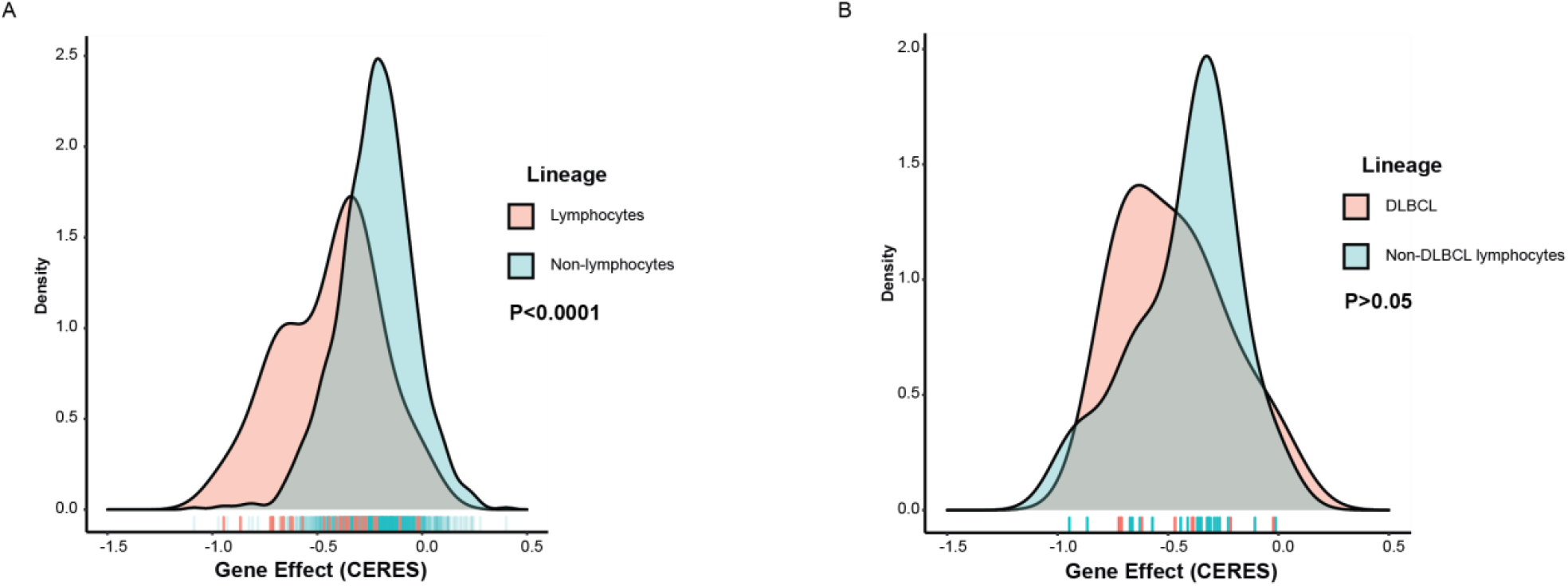
SLC25A1 dependency in cancer cell lines. Related to Figure 7. (A) Distribution of dependency for SLC25A1 in cancer cell lines of lymphocyte lineage versus cancer cell lines from all other lineages. (B) Distribution of dependency for SLC25A1 in DLBCL versus other lymphocyte lineage subtypes. Tick marks indicate individual cell lines. Dependency data downloaded from DepMap release 21Q2.

