## Supplementary material for "Mitochondrial pyruvate supports lymphoma proliferation by fueling a non-canonical glutamine metabolism pathway": Methods

STAR Methods:

**RESOURCE AVAILABILITY**

***Lead Contact***

***Materials Availability***

All unique/stable reagents generated within this study are available from the lead contact upon request without restriction.

***Data and Code Availability***

Code available at Github (https://github.com/alexbott/Wei_2021).

**EXPERIMENTAL MODEL AND SUBJECT DETAILS**

**DLBCL cell lines**

DLBCL cell lines used in this study (Pfeiffer, Toledo, OCI-Ly4, Karpas 422, U2932, OCI-Ly1, OCI-Ly7, SU-DHL-4, SU-DHL-6, and HBL-1) have been previously described (Caro et al., 2012; Norberg et al., 2017). All DLBCL cell lines were grown in RPMI 1640 medium with 2 g/L glucose, 0.3 g/L glutamine (Thermo Fisher 11875) supplemented with 10% FBS (Sigma, F0926) and 1% penicillin/streptomycin (HyClone) at 37 °C in a humidified atmosphere containing 5% CO_2_.

**Stable GDH and GPT2 knockdown cell lines**

HEK293T cells were transiently transfected with pLKO.1 shGDH-1 (Sigma Aldrich TRCN0000028600), shGDH-2 (Sigma Aldrich TRCN0000028611), shGPT2 (Sigma Aldrich TRCN0000035028) or scramble shRNA control (Addgene 8453) along with the lentiviral packaging plasmids pRSV-Rev, pMDLg/pRRE and pMD2.G using Lipofectamine 2000 transfection reagent. Forty-eight hours after transfection, viral supernatant was collected, filtered through a 0.45 µm polyethersulfone membrane, and stored at 4 °C. Ten µg/mL Polybrene (EMD Millipore, TR-1003-G) was added and a 1:1 mixture of viral supernatant and fresh growth medium (RPMI 1640 + 10% FBS) was applied directly to Pfeiffer cells, which were then incubated for 16 hours at 37 °C in a humidified incubator with 5% CO_2_. Viral media was discarded and replaced with fresh growth media and cells were allowed to recover and expand for 48 hours. After the recovery period, stably infected cells were selected with 1 µg/mL puromycin for 1 week. Knockdown of GDH and GPT2 were confirmed via immunoblotting as described below.

**METHOD DETAILS**

**SDS-PAGE and immunoblotting**

Whole cell lysates (WCLs) were prepared by scraping cells directly into RIPA buffer (50mM Tris-HCl, 1% NP-40, 0.5% Sodium Deoxycholate, 0.1% SDS, 150 mM NaCl, 2 mM EDTA) supplemented with protease and phosphatase inhibitors (Sigma Aldrich P8340, Roche Molecular 04906845001), incubated on ice for 45 minutes with vortexing every 5 minutes, and then spun at 16,000 × g for 10 minutes at 4 °C to remove insoluble material. WCL was normalized for total protein content via BCA Assay (Thermo Scientific 23225). Samples were resolved on SDS-PAGE gels and transferred to nitrocellulose membranes. Immunoblotting was performed using the indicated primary antibodies, which are listed in the key resources table according to the manufacturers’ recommendations, and analyzed on a LICOR Odyssey CLx.

**Cell growth and proliferation assays**

Cells from suspension culture were removed from flasks, mixed at a 1:1 ratio with 0.4% trypan blue solution (Sigma Aldrich T8154), and counted using an automated cell counter (Bio-rad 1450102). Cells from matrigel culture were scraped off plates with matrigel and media, spun at 600 × g for 10 minutes at 4 °C to remove media and matrigel. Cell pellets were resuspended in PBS, mixed at a 1:1 ratio with 0.4% trypan blue solution (Sigma Aldrich T8154), and counted using an automated cell counter (Bio-rad 1450102)

**Matrigel cell culture**

Cells were resuspended in ice-cold fresh RPMI 1640 growth medium at 2× concentration, then mixed at a 1:1 ratio with growth factor reduced matrigel (Corning 356231). 200 μL of mix was spotted into seven small drops in one well of a 6-well plate. After incubating for 30 minutes at 37 °C and 5% CO_2_ to allow the mix to polymerize, 3 mL of warm RPMI 1640 growth medium was added. Media was changed every 48 hours.

**^13^C-Glucose Tracing Experiments**

For suspension-culture tracing, 2 million cells were resuspended in 3 mL ^13^C-Glucose tracing media: glucose free RPMI 1640 (Thermo Fisher 11879020) supplemented with 11.11mM [U-^13^C]glucose, and 10% dialyzed FBS. For matrigel-culture tracing, 1.5 million cells were plated with matrigel then incubated with 3 mL growth media for 48 hours before the tracing experiment. To start experiment, media was changed to tracing media. Cells were harvested 30 minutes, 1 hour, 2 hours, and 4 hours later by centrifuge (scraped then centrifuge for cells from matrigel) and then quenched with 800 μL 80:20 methanol:water. Methanol lysates were then subjected to three rapid freeze-thaw cycles and then spun at 16,000 × g for 10 minutes at 4 °C. The supernatants were evaporated using a SpeedVac concentrator.

**^13^C-Glutamine Tracing Experiments**

For suspension-culture tracing, 2 million cells were resuspended in 3 mL ^13^C-Glutamine tracing meda: glutamine free RPMI 1640 (Thermo Fisher 21870076) supplemented with 2.05 mM [U-^13^C]glutamine, and 10% dialyzed FBS. For matrigel-culture tracing, 1.5 million cells were plated with matrigel then incubated with 3 mL growth media 48 hours before tracing experiment. To start experiment, media was changed to tracing media. Cells were harvested 30 minutes, and 2 hours later by centrifuge (scrape then centrifuge for cells from matrigel) and then quenched with 800 μL 80:20 methanol:water. Methanol lysates were then subjected to three rapid freeze-thaw cycles and then spun at 16,000 × g for 10 minutes at 4 °C. The supernatants were evaporated using a SpeedVac concentrator.

**^15^N-Glutamine Tracing Experiments**

For suspension-culture tracing, 2 million cells were resuspended in 3 mL [Alpha-^15^N]glutamine tracing meda: glutamine free RPMI 1640 (Thermo Fisher 21870076) supplemented with 2.05 mM [Alpha-^15^N]glutamine, and 10% dialyzed FBS. For matrigel-culture tracing, 1.5 million cells were plated with matrigel then incubated with 3 mL growth meda 48 hours before tracing experiment. To start experiment, media was changed to tracing media. Cells were harvested 4 hours later by centrifuge (scrape then centrifuge for cells from matrigel) and then quenched with 800 μL 80:20 methanol:water. Methanol lysates were then subjected to three rapid freeze-thaw cycles and then spun at 16,000 × g for 10 minutes at 4 °C. The supernatants were evaporated using a SpeedVac concentrator.

**Gas Chromatography Mass Spectrometry (GCMS) Derivatization**

The supernatants from tracing experiments were evaporated using a SpeedVac. Dried metabolites were re-suspended in 30 μL anhydrous pyridine with 10 mg/mL methoxyamine hydrochloride and incubated at room temperature overnight. The following morning, the samples were heated at 70 °C for 10–15 minutes and then centrifuged at 16,000 × g for 10 minutes. The supernatant was transferred to a pre-prepared GC/MS autoinjector vial containing 70 μL N-(tert-butyldimethylsilyl)-N-methyltrifluoroacetamide (MTBSTFA) derivitization reagent. The samples were incubated at 70 °C for 1 hour following which aliquots of 1 μL were injected for analysis. Samples were analyzed using either an Agilent 6890 or 7890 gas chromotograph coupled to an Agilent 5973N or 5975C Mass Selective Detector, respectively. The observed distributions of mass isotopologues were corrected for natural abundance.

**Steady State Metabolomics Experiments**

For matrigel samples, 2 million U2932 cells were plated with matrigel then incubated with 3 mL fresh growth medium in each plate. In parallel, 2 million same passaged cells were resuspended in 3 mL fresh growth medium in each flask. Cells were harvested 4 hours, 8 hour, 12 hours, and 24 hours later by centrifuge (scrape then centrifuge for cells from matrigel) and then quenched with 800 μL 80:20 methanol:water. Methanol lysates were then subjected to three rapid freeze-thaw cycle and then spun at 16,000 × g for 10 minutes at 4 °C. The supernatants were evaporated using a SpeedVac concentrator.

**Liquid Chromatography Mass Spectrometry (LCMS)**

The metabolite supernatants were evaporated using a SpeedVac. Dried metabolites were reconstituted in 100 μL of 0.03% formic acid in analytical-grade water, vortexed and centrifuged to remove insoluble material. The supernatant was collected and subjected to screening metabolomics analysis as described on an AB SCIEX QTRAP 5500 liquid chromatography/triple quadrupole mass spectrometer (Applied Biosystems SCIEX) (Kim et al., 2017). The injection volume was 20 μL. Chromatogram review and peak area integration were performed using MultiQuant (version 2.1, Applied Biosystems SCIEX). The peak area for each detected metabolite was normalized against the total ion count of that sample.

**Xenografts**

Ten million HBL1 Scramble or MPC knockdown cells were subcutaneously injected to the flank of the NRG mice in 1:1 PBS/Matrigel mix. Tumor sizes were measured at indicated times by caliper.

**Microarray Data Analysis**

Data were obtained from GSE10846 using the package GEOquery (version 2.60.0). BCR and OxPhos classifications were assigned according to Caro et al. (2012). Differential expression between the two groups was determined using limma (version 3.48.1).

**QUANTIFICATION AND STATISTICAL ANALYSIS**

For tracing and steady-state metabolomics, cell growth assays, and cell viability analysis, statistically significant differences were determined using GraphPad Prism 8. Data were analyzed by one-way Anova followed by Dunnett’s multiple comparison test. A p-value less than 0.05 was considered to be statistically significant. For microarray data analysis, statistical analysis was performed in R version 4.0.3.

**KEY RESOURCES TABLE**

| REAGENT or RESOURCE | SOURCE | IDENTIFIER |
| --- | --- | --- |
| Antibodies | | |
| MPC1, Rabbit monoclonal | Cell Signaling | Cat# 14462;  RRID:AB_2773729 |
| MPC2, Rabbit monoclonal | Cell Signaling | Cat# 46141;  RRID:AB_2799295 |
| VDAC, Rabbit monoclonal | Cell Signaling | Cat# 4866;  RRID:AB_2272627 |
| GDH, Rabbit monoclonal | Cell Signaling | Cat# 12793;  RRID:AB_2750880 |
| GPT2, Rabbit polyclonal | Sigma Aldrich | Cat# HPA051514;  RRID:AB_2681516 |
| α-Tubulin, Mouse monoclonal | Cell Signaling | Cat# 3873;  RRID:AB_1904178 |
| Bacterial and Virus Strains | | |
| pLKO.1 | Addgene | Cat# 8453 |
| Biological Samples | | |
| Chemicals, Peptides, and Recombinant Proteins | | |
| D-[U-^13^C]glucose | Cambridge Isotopes | Cat# CLM-1396 |
| L-[U-^13^C]glutamine | Cambridge Isotopes | Cat# CLM-1822 |
| L-[alpha-^15^N]glutamine | Cambridge Isotopes | Cat# NLM-1016 |
| Lipofectamine 2000 transfection reagent | Invitrogen | Cat# 11668019 |
| Polybrene | EMD Millipore | Cat# TR-1003-G |
| UK-5099 | Sigma Aldrich | Cat# PZ0160 |
| CB-839 | Sigma Aldrich | Cat# 5337170001 |
| Dimethyl-α-ketoglutarate | Sigma Aldrich | Cat# 349631 |
| Ammonium chloride (NH_4_Cl) | Sigma Aldrich | Cat# A9434 |
| Matrigel (growth factor reduced) | Corning | Cat# 356231 |
| Critical Commercial Assays | | |
| Pierce BCA Assay | Thermo | Cat# 23225 |
| 0.4% trypan blue solution | Sigma Aldrich | Cat# T8154 |
| Deposited Data | | |
| Experimental Models: Cell Lines | | |
| Human: HEK293T cell line | ATCC | Cat# CRL-11268; RRID:CVCL_1926 |
| Human: Pfeiffer cell line | Caro et al., 2012 | RRID:CVCL_3326 |
| Human: Toledo cell line | Caro et al., 2012 | RRID:CVCL_3611 |
| Human: OCI-Ly4 cell line | Caro et al., 2012 | RRID:CVCL_8801 |
| Human: Karpas 422 cell line | Caro et al., 2012 | RRID:CVCL_1325 |
| Human: U2932 cell line | Caro et al., 2012 | RRID:CVCL_1896 |
| Human: OCI-Ly1 cell line | Caro et al., 2012 | RRID:CVCL_1879 |
| Human: OCI-Ly7 cell line | Caro et al., 2012 | RRID:CVCL_1881 |
| Human: SU-DHL-4 cell line | Caro et al., 2012 | RRID:CVCL_0539 |
| Human: SU-DHL-6 cell line | Caro et al., 2012 | RRID:CVCL_2206 |
| Human: HBL-1 cell line | Caro et al., 2012 | RRID:CVCL_4213 |
| Recombinant DNA | | |
| pLKO.1 scramble (scramble shRNA) | Addgene | Cat# 8453 |
| Human shMPC1 | Sigma Aldrich | Cat# NM_016098;  TRCN0000005485 |
| Human shMPC2 | Sigma Aldrich | Cat# NM_015415;  TRCN0000278229 |
| Human shGDH-1 | Sigma Aldrich | Cat# NM_005271; TRCN0000028600 |
| Human shGDH-2 | Sigma Aldrich | Cat# NM_005271; TRCN0000028611 |
| Human shGPT2-1 | Sigma Aldrich | Cat# NM_133443; TRCN0000035028 |
| Human shGPT2-2 | Sigma Aldrich | Cat# NM_133443;  TRCN0000035024 |
| CRISPR Target Sequences | | |
| MPC1: AGGTTTACTGGGTTAATTGA, TAGATGCGCTTTAGCAGTTG | | |
| MPC2: AGGGATCGTTGGCAGCCGGG, TGGGTTGGAGTCGTGCGTAA | | |
| Software and Algorithms | | |
| Prism 9 |  |  |
| R Project for Statistical Computing | R Core Team, 2020 | RRID:SCR_001905 |
| pheatmap | Kolde, 2019 | RRID:SCR_016418 |
| ggplot2 | Wickham, 2016 | RRID:SCR_014601 |
| limma | Ritchie et al., 2015 | RRID:SCR_010943 |
